# Image Processing and Quality Control for the first 10,000 Brain Imaging Datasets from UK Biobank

**DOI:** 10.1101/130385

**Authors:** Fidel Alfaro-Almagro, Mark Jenkinson, Neal K. Bangerter, Jesper L. R. Andersson, Ludovica Griffanti, Gwenaëlle Douaud, Stamatios N. Sotiropoulos, Saad Jbabdi, Moises Hernandez-Fernandez, Emmanuel Vallee, Diego Vidaurre, Matthew Webster, Paul McCarthy, Christopher Rorden, Alessandro Daducci, Daniel C. Alexander, Hui Zhang, Iulius Dragonu, Paul M. Matthews, Karla L. Miller, Stephen M. Smith

## Abstract

UK Biobank is a large-scale prospective epidemiological study with all data accessible to researchers worldwide. It is currently in the process of bringing back 100,000 of the original participants for brain, heart and body MRI, carotid ultrasound and low-dose bone/fat x-ray. The brain imaging component covers 6 modalities (T1, T2 FLAIR, susceptibility weighted MRI, Resting fMRI, Task fMRI and Diffusion MRI). Raw and processed data from the first 10,000 imaged subjects has recently been released for general research access. To help convert this data into useful summary information we have developed an automated processing and QC (Quality Control) pipeline that is available for use by other researchers. In this paper we describe the pipeline in detail, following a brief overview of UK Biobank brain imaging and the acquisition protocol. We also describe several quantitative investigations carried out as part of the development of both the imaging protocol and the processing pipeline.

## 1. Introduction

### 1.1. Biobank Project

UK Biobank is a prospective cohort study of over 500,000 individuals from across the United Kingdom (Sudlow et al. (2015)). Voluntary participants, aged between 40 and 69 when recruited in 2006–2010, were invited to one of 22 centres across the UK. Blood, urine and saliva samples were collected, samples for genetic analysis and physical measurements were taken, and each individual answered an extensive questionnaire focused on aspects of health and lifestyle. This valuable data resource will provide insight into how the health of the UK population develops over many years and will enable researchers to improve the diagnosis and treatment of common diseases which will inevitably occur in sub-groups of the population.

In 2014, UK Biobank began the process of inviting back 100,000 of the original volunteers for brain, heart and body imaging. Imaging data for 10,000 volunteers has already been processed and made available for further research (Biobank, 2016, 2017). Due to the wide scope (in population size and number of imaging modalities) of the brain imaging acquisitions, UK Biobank will become an important resource for the research community. The included modalities are T1 (Section 3.2), T2 FLAIR (Section 3.3), swMRI (susceptibility-weighted MRI, Section 3.4), dMRI (diffusion MRI, Section 3.6), rfMRI (resting-state functional MRI, Section 3.7) and tfMRI (task fMRI, Section 3.8). It is hoped that ASL (arterial spin labelling, a measure of blood flow in the brain) can be added to the protocol in the near future. Section 2.1 briefly describes the acquisition protocol and the acquired modalities.

Other large-scale imaging projects include the Maastricht Study with 10,000 subjects (Schram et al. (2014)), the German National Cohort with 30,000 subjects (GNC (2014)) and the Rhineland Study with 30,000 subjects (Breteler et al., 2014).

The full potential of UK Biobank stems from its access to NHS (UK National Health Service) records; participants have agreed for their future health outcome information to be linked into the UK Biobank database. Thanks to this, researchers can aim to find novel biomarkers for early diagnosis of different diseases. As described in Sudlow et al. (2015), 1,800 of the imaged participants are expected to develop Alzheimer’s disease by 2022, rising to 6,000 by 2027 (diabetes: 8,000 rising to 14,000; stroke: 1,800 to 4,000; Parkinson’s: 1,200 to 2,800).

Greater detail regarding the background to the UK Biobank brain imaging study is given in Miller et al. (2016), a resource paper that also provides an introduction to the types of biological information that can be extracted from the different brain imaging modalities, as well as example results from early analyses from the first 5,000 subjects’ data, and detailed discussions of the challenges of analysing and interpreting such datasets. In contrast, here we focus more specifically on the methodological advances and implementation details in the image processing and QC pipeline that we have developed for UK Biobank; we describe here version 1.0 of “FBP” (FMRIB’s Biobank Pipeline), used to generate the first 10,000 subjects’ processed and released data, as of January 2017. We also include several investigations relating to acquisition and analysis pipeline decisions (such as B0 shimming approaches and registration method for dMRI data), with analyses designed to evaluate relative merits of different approaches.

### 1.2. Automated Pipeline

Brain imaging data is not immediately usable for extracting biologically-meaningful information in its raw form. It needs to be processed using a wide variety of software tools, optimised and combined together into a processing pipeline.

There are many excellent examples of processing pipelines in the literature. Tools such as those described in Wei et al. (2002) (segmentation), Cook et al. (2006) (diffusion) or Song et al. (2011) (resting-state), are specific solutions for the processing of specific image modalities. However, for the UK Biobank we need a robust pipeline that can process and integrate across many different functional and structural modalities. The minimal preprocessing pipelines developed by the Human Connectome Project (HCP) (Glasser et al., 2013) are good examples of pipelines optimised for the specific needs of a project with very special characteristics. Following the same philosophy, we have developed a fully automated processing pipeline primarily based around FSL software (Jenkinson et al., 2012). In Section 3 we describe the pipeline in detail, as well as the choices we have made in its development.

### 1.3. Imaging-Derived Phenotypes and Quality Control

A goal of UK Biobank is to identify new relationships between different phenotypic features and human diseases, in the hope that they may be used as biomarkers for early diagnosis. Therefore, in addition to preprocessing the brain imaging data (e.g., aligning modalities and removing artefacts), our pipeline also generates approximately 4350 “imaging-derived phenotypes” (IDPs). These IDPs include summary measures such as subcortical structure volumes, microstructural measures in major tracts (DTI and NODDI measures (Zhang et al., 2012)), and structural/functional connectivity metrics. IDPs for the first 10,000 subjects have been publicly released^1^ and can be used in combination with other non-imaging data from Biobank to identify disease risk factors and biomarkers. Each IDP is presented as a separate data field within the UK Biobank showcase (except for the rfMRI netmats and node amplitudes, which are saved en masse within “bulk” variables in the database). A detailed description of all IDPs is available in Section S3 of the supplementary material, and the complete list is given online^2^

Quality Control (QC) is a very important issue in brain imaging, as poorly executed application (or lack thereof) can compromise the trustworthiness of a study (Bennett and Miller, 2010). This topic has been explored in the literature, although very often research is mostly focused on Quality Assurance (QA) rather than QC. QA is focused on avoiding the occurrence of problems by improving a process while QC is focused on finding possible problems in the output of that process. For example, the Function Biomedical Informatics Research Network (FBIRN) set of recommendations (Glover et al., 2012) is solely focused on QA. In a similar vein, Friedman and Glover (2006) explore an interesting set of quality metrics, but they focus on stability, signal-to-noise ratio (SNR), drift, and other hardware performance issues related to MR scanners, not specifically on the type of artefacts^3^ that can be found in MR imaging even when complying with the best QA policies.

The scale of a typical brain imaging study to date (up to 100 subjects) allows researchers to largely perform QC manually, by visually inspecting the data at each step in the processing pipeline. Even the HCP, while having a relatively large number of subjects (1200), relies primarily upon visual inspection for their T1 QC process (Marcus et al., 2013). However, the sheer quantity of imaging data which will be produced by UK Biobank (100,000 subjects) makes QC via visual inspection unfeasible. Thus, the development of reliable QC metrics to detect artefacts specific to the imaging acquisition and analysis process is essential.

Various imaging-related QC metrics have been proposed (e.g. Deshmukh and Bhosale (2010), Moorthy and Bovik (2010)) but, as they are not specific to MR or brain imaging they do not fulfill the needs of our study. More specific proposals can be found in other studies. Woodard and Carley-Spencer (2006) suggests a metric based on Natural Scene Statistics to detect noise in structural images. Following the same idea, Mortamet et al. (2009) suggests two metrics to detect noise (based on investigating the ratio of artefactual voxels relative to the background, or the noise distribution in the background). Those proposals, although extremely useful, refer to a very specific problem in structural images, and hence are unlikely to detect artefacts due to other sources, or in other modalities.

Similarly, there have been proposed metrics for dMRI (Hasan, 2007; Liu et al., 2010), fMRI (Greve et al., 2011; Power et al., 2012; Nichols, 2013; Afyouni and Nichols, 2017), and more recent multimodal approaches (Abe et al., 2015), but the extent to which these metrics may capture the majority of artefacts that could arise is yet to be proven.

Finally, PCP QAP (Preprocessed Connectors Project Quality Assessment Protocol)^4^ (Craddock et al., 2016) is a concerted effort to create a unified platform for Quality Control that attempts to incorporate different QC pipelines and their associated metrics. The QC tools to come out of our work are designed to be easily integrated into MRIQC (Esteban et al., 2016, 2017), a project affiliated to PCPQAP.

Due to the uncertainty about the suitability of the QC metrics discussed above to successfully assess image quality objectively and to detect the majority of brain artefacts, we propose a machine learning approach to automatically identify problematic images based on a wide range of imaging derived metrics. For this study, the generated IDPs are combined with a collection of additional QC-specific metrics (e.g., measures of asymmetry, normalised intensity per subcortical structure, metrics to detect alignment that classifies images as usable or non-usable). This has been developed for, and tested on, T1 images, and will be further developed in future work so that it can be applied to all modalities. Section 4 further describes the QC-specific metrics and the machine learning system. The main reason for our heavy focus to date on QC for the T1 data is that the entire processing pipeline (for all modalities) relies heavily on having a usable T1 structural image.

## 2. Data acquisition

### 2.1. Acquisition overview

UK Biobank brain imaging has been described as a resource in Miller et al. (2016), where the reader can find a number of examples describing how it can be accessed and used for research. The data is available online^5^.

Three dedicated imaging centres are equipped with identical scanners (3T Siemens Skyra, software VD13) for brain imaging scanning using the standard Siemens 32-channel receive head coil. As shown in multiple studies (Focke et al., 2011; Chen et al., 2014), having the same scanner and protocol is advantageous for a multi-site study.

The scanners operate 7 days per week, each scanning 18 subjects per day. These acquisition rates demand an extremely time-efficient protocol - each minute of scanning (added into the protocol) in effect costs £1 million, so scanning time has been a key criterion in the protocol development. The total scanning time has been established as 31 minutes (plus 5 minutes for subject adjustment, shimming, etc). The optimised order of acquisition is: (1) T1, (2) resting fMRI, (3) task fMRI, (4) T2 FLAIR, (5) dMRI, (6) swMRI (Biobank, 2014).

During the acquisition process, if a significant artefact is observed while scanning, and there is enough time to restart the sequence, then the acquisition may be repeated. The most common reason for these artefacts is excessive head movement.

Likewise, as mentioned in Miller et al. (2016), in case of a possible health-related incidental finding noted by the radiographer during the imaging process, a formal radiologist review is undertaken and, if it is confirmed as potentially serious, feedback is given to the participant and their doctor.

To date, our processing pipeline has been applied to 10,129 UK Biobank volunteers scanned from 2014 to 2016 at just the first imaging centre. The main acquisition parameters for each modality are listed in Table 1.

A usable T1-weighted image was obtained for 98% of these ∼10,000 subjects (the same figure that was reported in Miller et al. (2016) for the initial ∼6000 subjects, confirming the stability of the acquisition process). This is an important point, as a T1 image is a crucial part of the processing pipeline, being essential for making full use of data from the other modalities. At this stage, an image was deemed unusable only after careful manual quality checks across all modalities^6^. A set of manual classification categories, specific to each modality, were assigned to every image; these categories are described in section 4.1. Data for all modalities were acquired and usable in 8211 subjects (81.06%).

Throughout the development of the Biobank imaging procedures, the imaging acquisition has been divided into different protocol phases, with each phase corresponding to (generally minor early) adjustments in the acquisition protocol. The original aim in the UK Biobank brain imaging component was to maintain a fixed acquisition protocol during the 5-6 years that the scanning will need or, at least, to have maximum compatibility with eventual changes. There have been a number of protocol optimisations, particularly during the very earliest scanning periods (and hence covering a relatively small number of the earliest participants). There also have been a number of minor software upgrades (for obvious reasons the imaging study seeks to minimise any major hardware or software changes). Detailed descriptions of every protocol change (even the most minor), along with thorough investigations of the effects of these on the resulting data, will be the subject of a future paper. In Section S1 of supplementary material we describe one protocol change for which we carried out specific custom analyses to help verify the protocol change decision (relating to field shimming).

**Table 1.** UK Biobank brain MRI protocol (31 minutes total scan time).

| Modality | Duration | Voxel, Matrix | Key Parameters | Timepoints |
| --- | --- | --- | --- | --- |
| T1 | 4:54 | $1 \times 1 \times 1\text{mm}$<br>$208 \times 256 \times 256$ | 3D MPRAGE, sagittal, R=2,<br>TI/TR=880/2000 ms | 1 |
| T2 FLAIR | 5:52 | $1.05 \times 1.0 \times 1.0\text{mm}$<br>$192 \times 256 \times 256$ | FLAIR, 3D SPACE, sagittal,<br>R=2, PF 7/8, fat sat,<br>TI/TR=1800/5000 ms, elliptical | 1 |
| swMRI | 2:34 | $0.8 \times 0.8 \times 3.0\text{mm}$<br>$256 \times 288 \times 48$ | 3D GRE, axial, R=2, PF 7/8<br>TE1/TE2/TR=9.4/20/27ms | 2 |
| dMRI | 7:08 | $2.0 \times 2.0 \times 2.0\text{mm}$<br>$104 \times 104 \times 72$ | MB=3, R=1, TE/TR=92/3600<br>ms, PF 6/8, fat sat, $b=0 \text{ s/mm}^2$<br>(5x + 3x phase-encoding<br>reversed), $b=1000 \text{ s/mm}^2$ (50x),<br>$b=2000 \text{ s/mm}^2$ (50x) | 105 + 6 (AP<br>+ PA) |
| rfMRI | 6:10 | $2.4 \times 2.4 \times 2.4\text{mm}$<br>$88 \times 88 \times 64$ | TE/TR=39/735 ms, MB=8,<br>R=1, flip angle $52^\circ$, fat sat | 490 |
| tfMRI | 4:13 | $2.4 \times 2.4 \times 2.4\text{mm}$<br>$88 \times 88 \times 64$ | Acquisition same as rfMRI. Task<br>is faces/shapes “emotion” task. | 332 |

R = in-plane acceleration factor, MB = multiband factor, PF=partial Fourier. All non-EPI scans are pre-scan normalised (on-scanner bias-field corrected). Gradient distortion correction is deselected on the scanner and applied in post-processing. SNR, TSNR and motion plots are shown in figures S10 and S11 of the supplementary material.

### 2.2. Acquired modalities overview

**T1-weighted structural imaging (“T1”, Mugler and Brookeman (1990))** provides information relating to volumes and morphology of brain tissues and structures. It is also critical for calculations of cross-subject and cross-modality alignments, needed in order to process all other brain modalities. Acquisition details are: 1mm isotropic resolution using a 3D MPRAGE acquisition. The superior-inferior field-of-view is large (256mm), at little cost, in order to include reasonable amounts of neck/mouth, as those areas hold interest for some researchers.

**T2-weighted FLAIR imaging (“T2 FLAIR”, Mugler (2014))** is a structural technique with contrast dominated by signal decay from interactions between water molecules (T2 relaxation times). T2 images depict alterations to tissue properties typically associated with certain pathology (e.g., white matter lesions). In this modality, signal from fluid (CSF) is suppressed and hence it appears dark (unlike either PD or T2w images).

After early piloting, a standard **T2/PD-weighted acquisition** was dropped due to a combination of factors such as overall value and timing practicalities. However the T2-weighted FLAIR image is still acquired, which is generally of good quality and which shows strong contrast for white matter hyperintensities.

**Susceptibility-weighted imaging (“swMRI”, Haacke et al. (2004); Duyn (2013))** is a structural tech-nique that is sensitive to “magnetic” tissue constituents (magnetic susceptibility). Data from one scan (including phase and magnitude images from two echo times) can be processed in multiple ways to reflect venous vasculature, hemosiderin microbleeds, quantitative susceptibility mapping or aspects of microstructure (e.g., iron, calcium and myelin). Voxel size for this modality is 0.8 *×* 0.8 *×* 3.0mm. Anisotropic voxels enhance contrast in the signal magnitude from sources of signal dephasing, such as iron in veins or microbleeds, but are less ideal for other susceptibility-based processing. Ultimately, however, the decision to acquire anisotropic voxels was motivated by the desire for whole brain coverage in the face of very limited scan time (2.5 minutes).

**Diffusion-weighted imaging (“dMRI”, Sotiropoulos et al. (2013a); Xu et al. (2013); Sotiropoulos et al. (2013b))** measures the ability of water molecules to move within their local tissue environment. Water diffusion is measured along a range of orientations, providing two types of useful information. Local (voxel-wise) estimates of diffusion properties reflect the integrity of microstructural tissue compartments (e.g., diffusion tensor and NODDI measures). Long-range estimates based on tract-tracing (tractography) reflect structural connectivity between pairs of brain regions. For the two diffusion-weighted shells, 50 diffusion-encoding directions were acquired (with all 100 directions being distinct). Acquisition phase-encoding direction for the dMRI data is AP (Anterior-to-Posterior). In order to generate appropriate fieldmaps to carry out geometric distortion correction for EPI images (See Section 3.5), a reverse (PA: Posterior-to-Anterior) phase-encoding dMRI is also acquired.

**Resting-state functional MRI (“rfMRI”)** measures changes in blood oxygenation associated with intrinsic brain activity (i.e., in the absence of an explicit task or sensory stimulus). It can generate valuable estimations of the apparent connectivity between pairs of brain regions, as reflected in the presence of spontaneous co-fluctuations in signal. As implemented in the CMRR multiband acquisition (Moeller et al., 2010), a separate “single-band reference scan” (SBRef) is acquired. This has the same geometry (including EPI distortion) as the timeseries data, but is not part of a low-TR timeseries, and hence, without the corresponding T1 saturation effects, has higher between-tissue contrast; this is used as the reference scan in head motion correction and alignment to other modalities. In cases where the SBRef image is missing, the alignments use an average image generated from the first 5 volumes from the fMRI timeseries and selecting the one that correlates most highly with the others (after co-alignment). This procedure is similar to the selection of the best b=0 image described in section 3.5.

**Task functional MRI (“tfMRI”)** uses the same measurement technique as resting-state fMRI, while the subject performs a particular task or experiences a sensory stimulus. The task used is the Hariri faces/shapes “emotion” task (Hariri et al., 2002; Barch et al., 2013), as implemented in the HCP, but with shorter overall duration and hence fewer total stimulus block repeats. The participants are presented with blocks of trials and asked to decide either which of two faces presented on the bottom of the screen match the face at the top of the screen, or which of two shapes presented at the bottom of the screen match the shape at the top of the screen. The faces have either angry or fearful expressions. This task was chosen to engage a range of sensory, motor and cognitive systems.

## 3. Automated processing pipelines

All processing was performed on a dedicated cluster composed of CPU servers (typical speed 2.66 GHz per core, average RAM 10 GB per core) as well as GPU servers for the GPU-optimised parts of the processing pipeline (NVidia Tesla K40/K80 GPUs for processing with Cuda 5.5^7^).

Image reconstruction from k-space is performed on the scanner using standard Siemens reconstructions, except for multiband EPI (reconstructed using custom CMRR code). Partial bias field correction in structural data is performed via the on-scanner “pre-scan normalise” option. No on-scanner gradient distortion correction is applied. After reconstruction and on-scanner preprocessing, image data is received (in DICOM format) by the pipeline. An overview of the processing pipeline can be seen in Figure 2.

The source code for the pipeline can be found online^8^.

**Figure 1:**
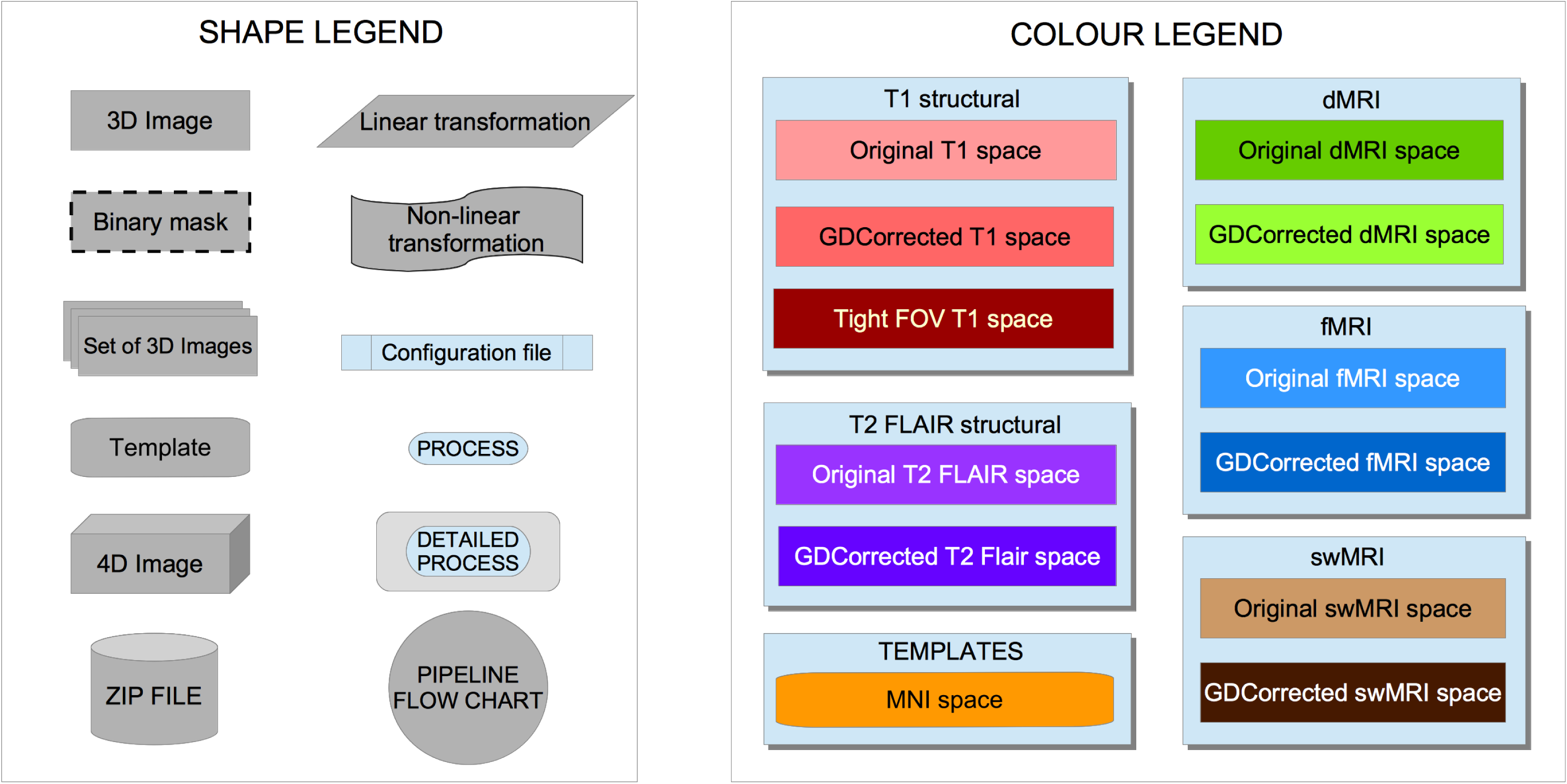
Flowcharts legend. All linear registrations shown in the pipeline are rigid body transformations (6 degrees of freedom) in intra-subject operations and affine transformations (12 degrees of freedom) when registering to MNI template.

**Figure 2:**
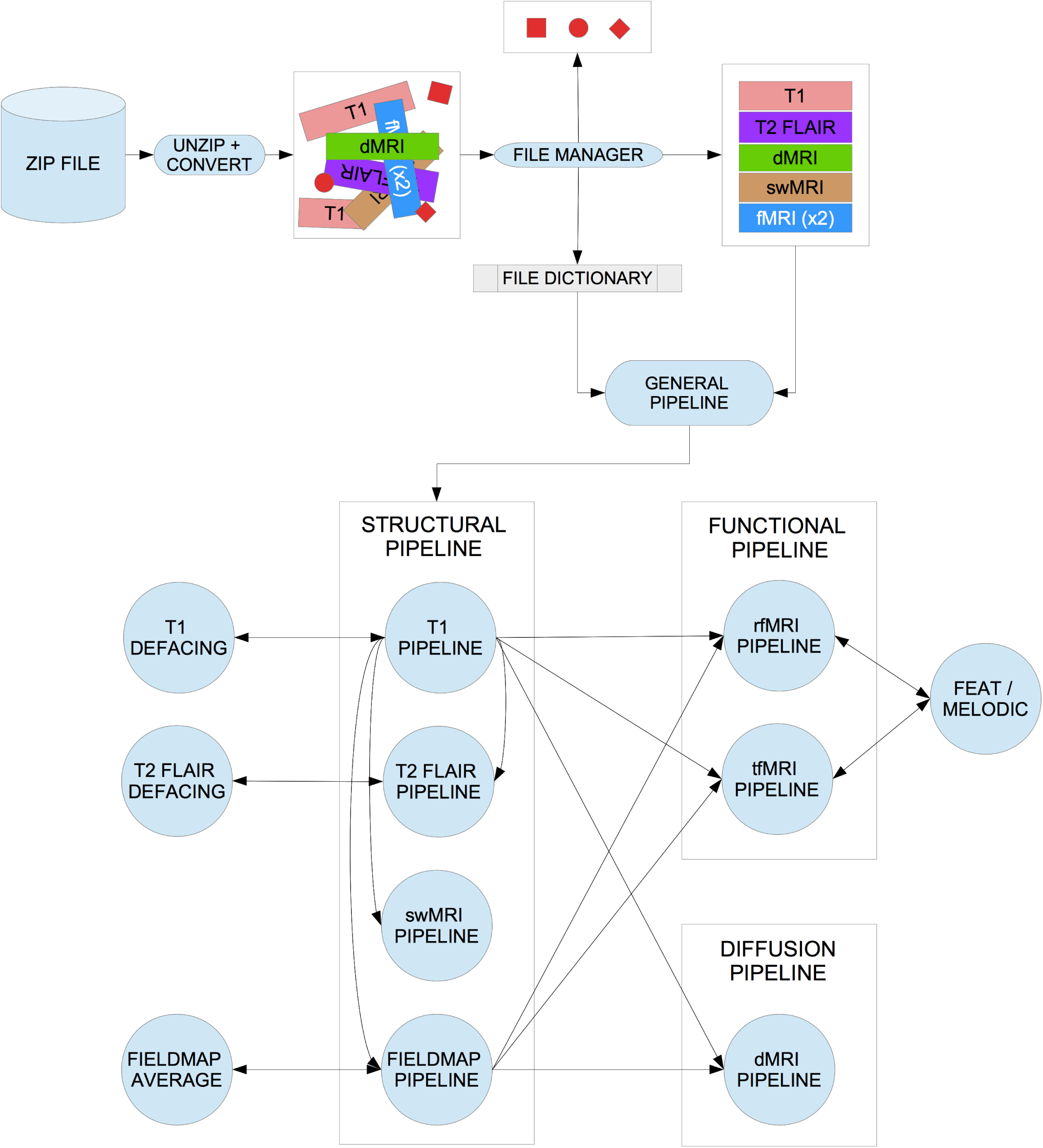
General flowchart for the pipeline.

### 3.1. Initial preprocessing steps

#### 3.1.1. Data conversion: DICOM to NIFTI

All DICOM image files are converted to NIFTI format using dcm2niix (Li et al., 2016). This tool also generates the diffusion-encoding b-value and b-vector files, as well as a JSON file for each NIFTI image with meta-information such as: acquisition date and time, echo time, repetition time, effective echo spacing, encoding direction, magnetic field strength, flip angle and normalisation by scanner.

Image data is available from UK Biobank in both DICOM and (separately) NIFTI formats. Both forms include the raw (non-processed) image-space data, the only differences being that the NIFTI-version T1/T2 structural images are defaced for subject anonymity (as described below), and the multi-coil (pre-combination) swMRI data is only available in the DICOM downloads. Researchers download individual zipfiles corresponding to one modality from one subject for one data format (DICOM or NIFTI).

The NIFTI versions are the recommended option, partly because for each modality a small number of simply and consistently named images are provided (e.g., “T1”, “rfMRI”), as opposed to thousands of separate DICOM files (with complex naming and somewhat variable conventions dictated by the scanner). Also, the NIFTI downloads, while overall only being 40% larger than pure-raw DICOM downloads, include not just the raw images, but also images output by the processing pipeline, for example after gradient distortion correction (for all modalities), and correction for eddy currents and head motion (dMRI data), and artefact removal (rfMRI data).

#### 3.1.2. Data organization

The directory tree structure of the processed data is available online^9^.

For each subject, the raw and processed imaging data files are automatically organised into subfolders according to the different modalities, with a uniform naming scheme for files and directories. To achieve this, we need to check all the files that have been converted from DICOM to NIFTI, find the ones that correspond to each modality (by matching a set of naming patterns), sort out the possible problems with the number of files found per modality (missing files or having more files than expected) and finally, renaming the result according to the naming scheme. To resolve problems regarding the number of found files, we developed the logical rules defined in Algorithms 1 to 4 in Section S2 of the supplementary material.

This file organisation can be converted to BIDS file-naming format (Gorgolewski et al., 2016) with a conversion script included in the pipeline.

After the main file organisation, a basic Quality Control (QC) tool is used to check if every raw dataset in each modality has the correct dimensions. When raw data has the wrong dimensions, is corrupted, missing or otherwise unusable, it is moved into a sub folder called ”unusable” (inside the given modality’s folder), and not processed any further (apart from defacing applied to the raw T1 and T2 FLAIR).

This “unusable” data is included in the NIFTI packages in Biobank database because some researchers may be interested in working with it, for example, to develop methods for detecting or even possibly correcting such data.

In the case of unusable T1 data, the raw imaging data for all other modalities is deemed unusable (because the pipeline cannot function without a usable T1). However, as with the T1 data, all such raw data is still available for NIFTI download, but without any processing applied.

**Figure 3:**
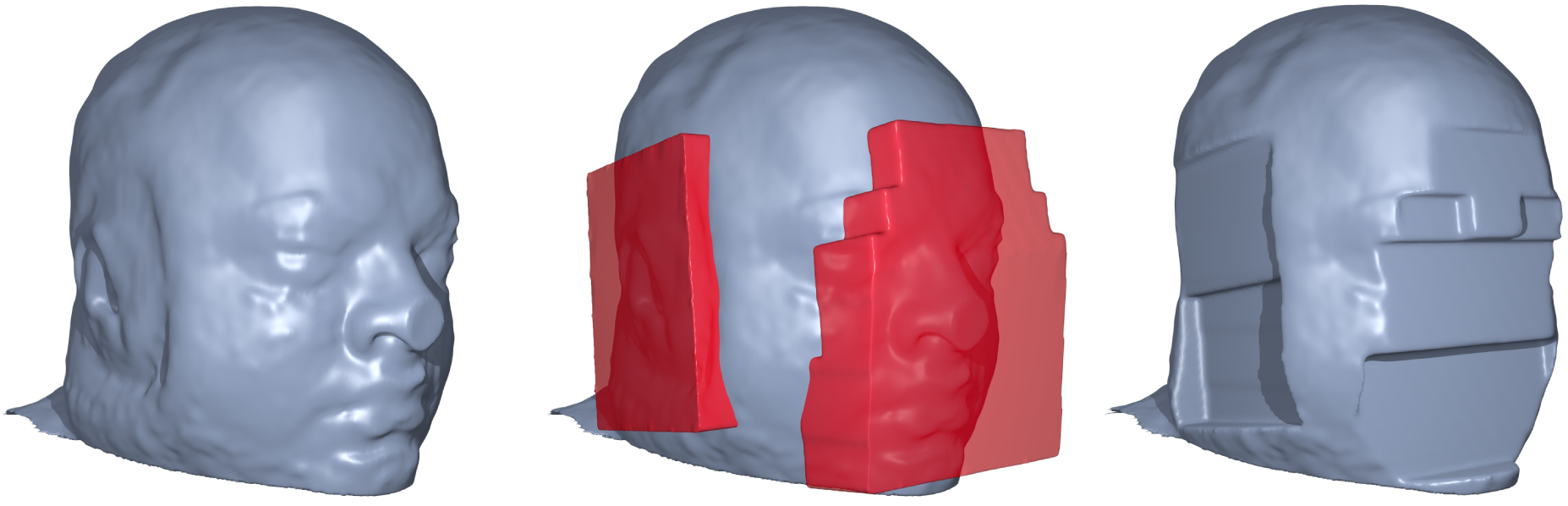
Example of defacing. *Left:* Original T1 volume (non-Biobank subject). *Center:* Applying defacing masks. *Right:* Defaced T1 volume.

#### 3.1.3. Image anonymisation

To protect study participant anonymity, the header data does not contain sensitive information such as name and any other information that could be used to identify the participant.

Furthermore, the high resolution structural images (T1 and T2) are automatically “defaced” by masking out voxels in the face and ear regions. This is accomplished by estimating a linear transformation between the original data and a standard co-ordinate system (an expanded MNI152 template space)^10^, and back-projecting a standard-space mask of the face and ear regions into the native data. This process is depicted in Figure 3. To ensure that this process did not unduly remove within-brain voxels, we calculated the overlap between non-defaced brain masks and the defacing masks. Overlap was present in only 0.5% of the subjects^11^ and, within these subjects, there was nearly no loss of brain voxels (worst case having 68 overlap voxels out of 1725983 brain voxels).

This defacing procedure is similar to common practice such as in the HCP. The raw, non-defaced DICOM T1 and T2 data is classified as “sensitive” by UK Biobank; researchers requiring raw DICOM non-defaced T1/T2 data should contact UK Biobank to discuss special access requirements.

#### 3.1.4. Gradient distortion correction

Full 3D gradient distortion correction (GDC) is not available on the scanner for EPI data (gradient distortions are only corrected within-plane for 2D EPI data), so GDC is entirely applied within the processing pipeline.

GDC is a necessary step, as shown in Figure 4. Tools developed by the FreeSurfer and HCP teams are used for applying the correction, available through the HCP GitHub account^12^. Running these tools also requires a proprietary Siemens data file that describes the gradient nonlinearities^13^.

### 3.2. T1 pipeline

T1 structural images are used as a reference for all other modalities. Processing performed on these images should avoid any kind of unnecessary smoothing or interpolation. Therefore, all linear and non-linear pipeline transformations are estimated, but not applied until as late as possible in the processing pipeline when all transforms can be combined and applied with a single interpolation.

**Figure 4:**
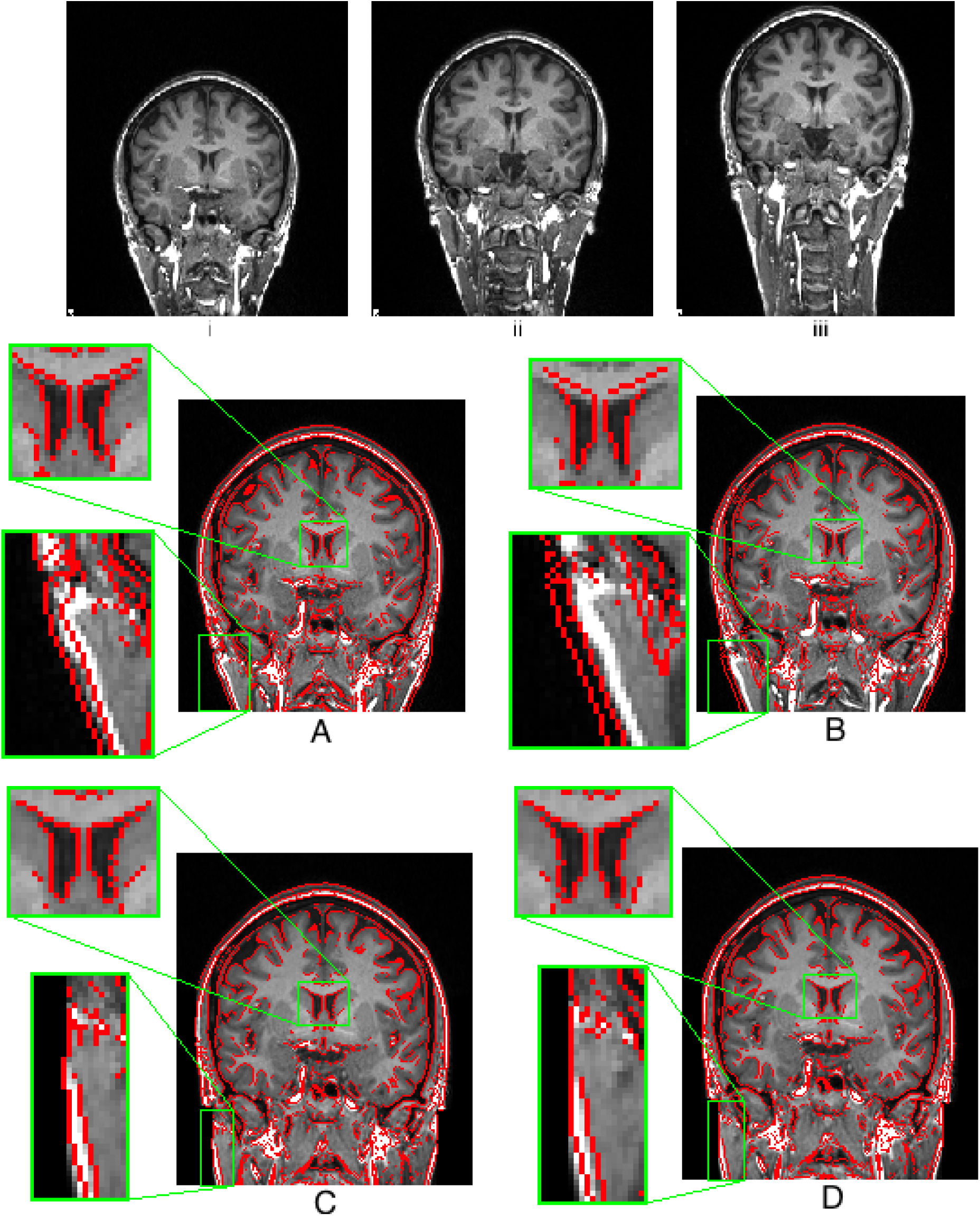
One volunteer was scanned for testing purposes at different positions on the scanning table, depicting the effect of gradient distortion (note the warping in the neck). Correction for gradient distortion (*C* and *D*) substantially improves alignment. *i:* Subject placed low in the scanner (Baseline). *ii:* Subject placed in the centre of the scanner. *iii:* Subject placed higher into the scanner. *A:* ii (red outline) linearly registered to i (background) (Cross correlation: 0.90). *B:* iii (red outline) linearly registered to i (background) (Cross correlation: 0.83). *C:* ii (red outline) linearly registered to i (background) after GDC (Cross correlation: 0.98). *D:* iii (red outline) linearly registered to i (background) after GDC (Cross correlation: 0.96). Regions highlighting the largest effect (i.e. improved alignment of the ventricle) of the GDC have been zoomed to a 3:1 scale.

As can be seen in Figure 5, the pipeline runs a GDC, cuts down the field of view (FOV), calculates a registration (linear and then non-linear) to the standard atlas, applies brain extraction, performs defacing and finally segments the brain into different tissues and subcortical structures.

In more detail: the pipeline first applies GDC for the original T1 image as described above. The FOV is then cut down to reduce the amount of non-brain tissue (primarily empty space above the head and tissue below the brain) to improve robustness and accuracy of subsequent registrations. Tools used to achieve this include BET (Brain Extraction Tool, Smith (2002)) and FLIRT (FMRIB’s Linear Image Registration Tool, Jenkinson and Smith (2001); Jenkinson et al. (2002))^14^, in conjunction with the MNI152 “nonlinear 6th generation” standard-space T1 template^15^. This results in a reduced-FOV T1 head image.

The next step is a non-linear registration to MNI152 space using FNIRT^16^ (FMRIB’s Nonlinear Image Registration Tool Andersson et al. (2007b,a)). This is a critical step in the pipeline, as many subsequent processing steps depend on accurate registration to standard space. We used the 1mm resolution version of MNI152 template as the reference space.

A particular issue for the non-linear registration was the fact that T1 images in UK Biobank have brighter internal carotid arteries than those the in MNI152 template. This affects the non-linear registration procedure, resulting in distortion in the temporal lobes. Hence, we created a custom reference brain mask to exclude this part of the image when estimating the transformation (see Figure S8 of the supplementary material).

All of the transformations estimated above (GDC, linear and non-linear transformations to MNI152) are then combined into one single non-linear transformation, which allows the original T1 to be transformed into MNI152 space (or vice versa) in a single step.

Using the inverse of the MNI152 alignment warp, a standard-space brain mask is transformed into the native T1 space and applied to the T1 image to generate a brain-extracted T1; this brain extraction replaces the earlier brain-extraction output created by BET (which was needed just for the initial registration stages). Similarly, as discussed avobe, a defacing mask (defined as a set of boxes removing eyes, nose, mouth and ears) is transformed into T1 space using the linear transformation and applied for anonymisation purposes. Only the linear transformation is used for defacing, as non-linear registration is performed on brain-extracted images, meaning that the warp field in out-of-brain regions is poorly conditioned. This process is shown in Figure 6.

Next, tissue-type segmentation is applied using FAST (FMRIB’s Automated Segmentation Tool Zhang et al. (2001)). This estimates discrete and probabilistic segmentations for CSF (cerebrospinal fluid), grey matter and white matter. As part of the segmentation, intensity bias is estimated, and so this step is also used to generate a fully bias-field-corrected version of the brain-extracted T1.

These data are then used to carry out a SIENAX^17^ analysis Smith et al. (2002). The external surface of the skull is estimated from the T1, and used to normalise brain tissue volumes for head size (compared with the MNI152 template). Volumes of different tissue types and total brain volume, both unnormalised and normalised for head size, are then generated.

Subcortical structures (shapes and volumes) are modelled using FIRST (FMRIB’s Integrated Registration and Segmentation Tool Patenaude et al. (2011)). The shape and volume outputs for 15 subcortical structures are generated and stored. A single summary image, with a distinct value coding for each structure is also generated.

From the T1 structural image, several global volume measures from SIENAX are reported as distinct IDPs, both normalised for overall head size as well as not normalised: total brain (grey + white matter) volume; total white matter volume, total grey matter volume, ventricular (non-peripheral) CSF (cerebrospinal fluid) volume; peripheral cortical grey matter volume. The overall volumetric head-size scaling factor is also recorded as an IDP. Several subcortical structures’ volumes from FIRST segmentation (not normalised for brain/head size, although that normalisation can be easily done, as the total brain volume is an IDP) are also reported, in general with separate IDPs for left and right, such as left thalamus and right thalamus. Total volume of grey matter (using the grey matter partial volume estimates from FAST) within 139 GM ROIs^18^ and total volume of white matter hyperintensities (WMHs) calculated with BIANCA (Griffanti et al., 2016) (using both T1 and T2 FLAIR) are also included as IDPs.

**Figure 5:**
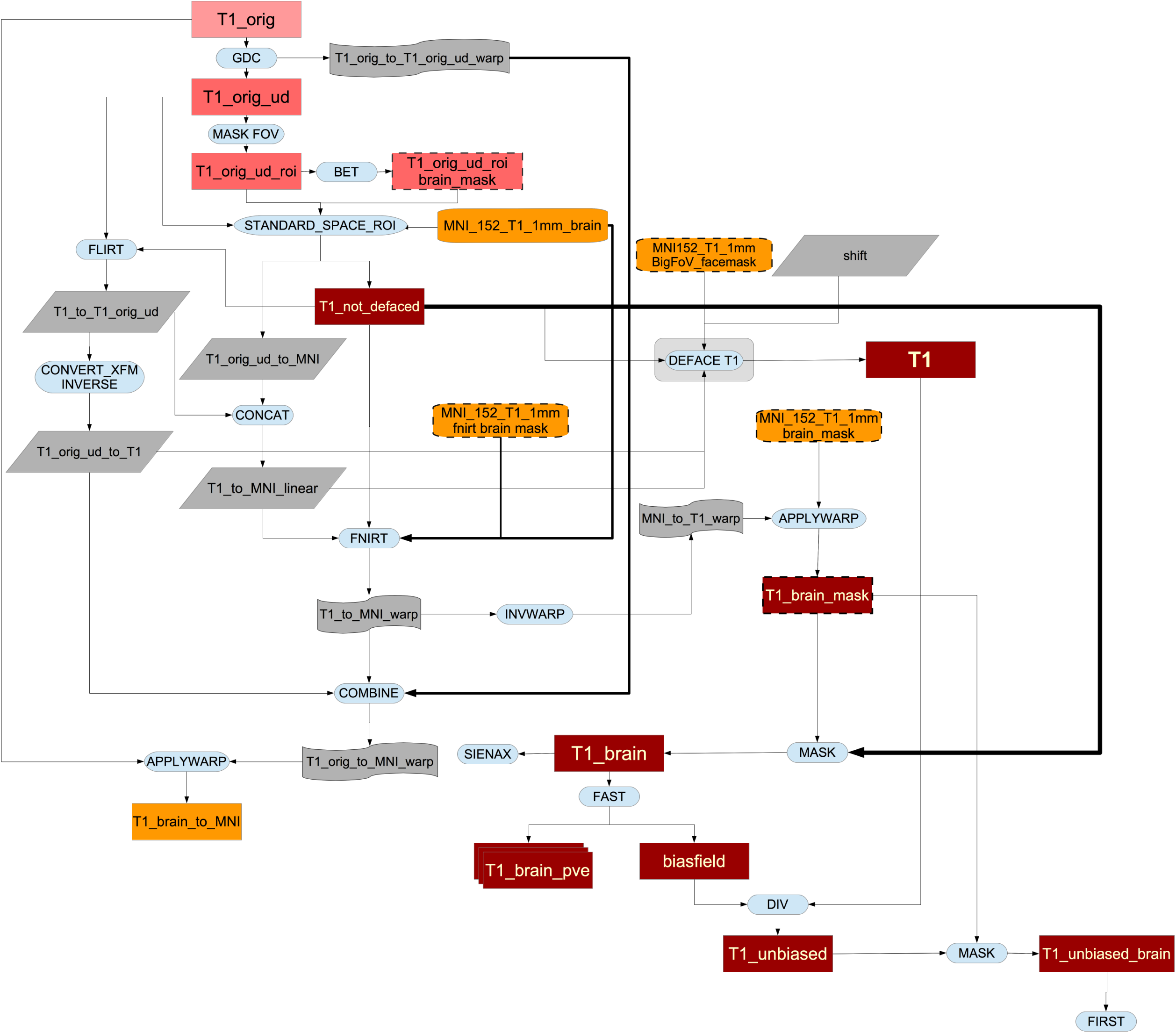
Flowchart for the T1 processing pipeline.

**Figure 6:**
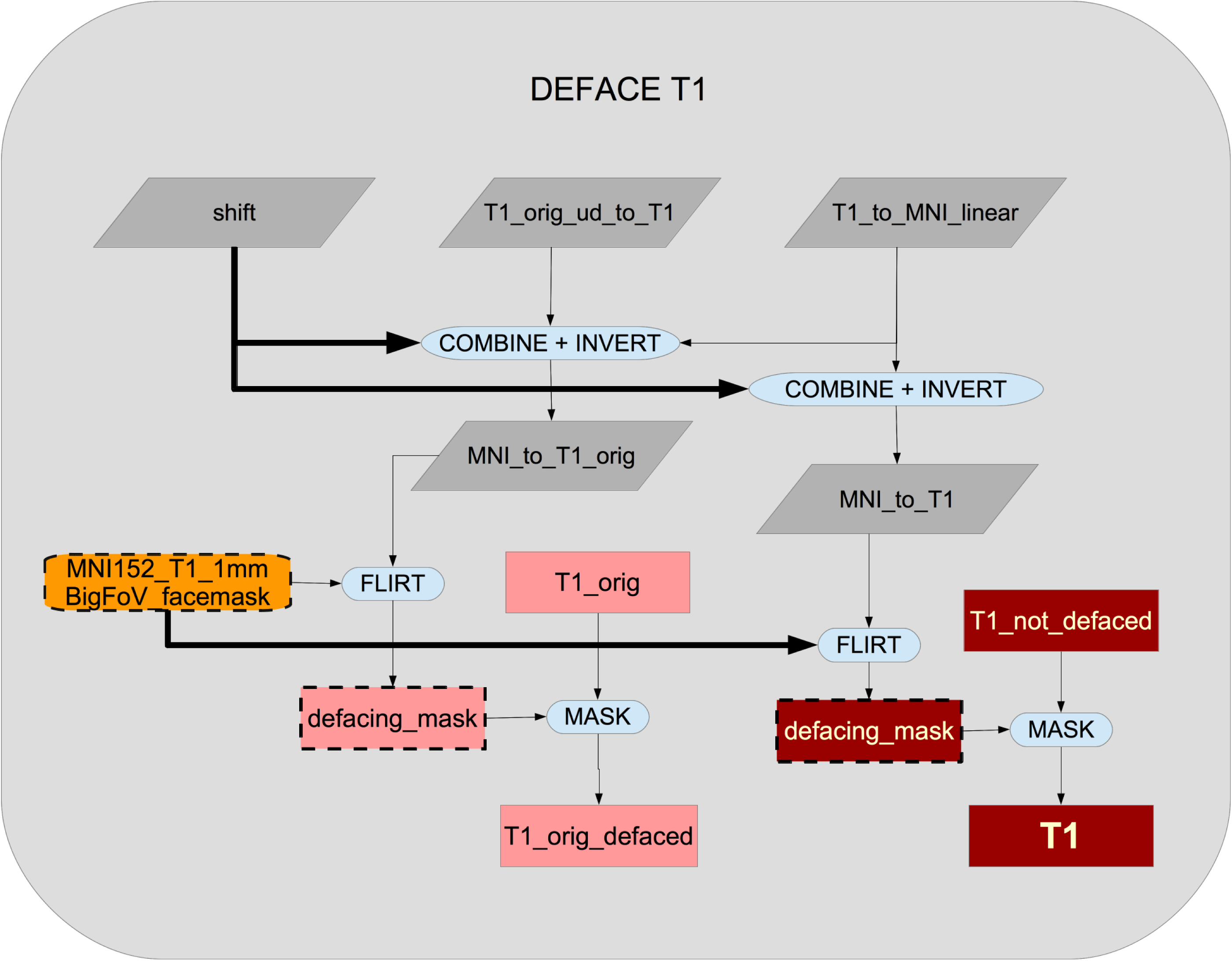
Flowchart for the defacing of the T1.

### 3.3. T2 FLAIR pipeline

The T2 FLAIR processing pipeline is very similar to the T1 pipeline, although we use the non-linear T1-to-MNI152 transformation to transform T2 FLAIR to MNI152 space. For this reason and to assist other combined analyses (e.g., white-matter hyperintensity segmentation), the T2 FLAIR image is first transformed to T1 space as described below.

As shown in Figure 7, the original T2 FLAIR image is first corrected for gradient distortions (GDC) and then a rigid-body (6 degrees of freedom) linear registration using FLIRT is applied to transform the corrected T2 FLAIR into corrected T1 space. After this step, the T1 brain and defacing masks are applied to the T2 FLAIR image (see Figure 8). Finally, an MNI152-version of T2 FLAIR is also generated using the previously calculated warp from T1 to MNI152.

As a last step, we use the estimated bias field previously calculated by FAST on the T1 image to correct the residual bias field inhomogeneities in the T2 FLAIR image. This approach performs as well as estimating the bias field directly from the T2 FLAIR that have not been removed by the on-scanner ”pre-scan normalise” processing. To validate this approach, we confirmed that the histograms for white matter intensity in 78 (randomly chosen) subjects improved, i.e., the distribution of WM intensity (as measured by the Inter Quartile Range [IQR]) was tightened in a similar way for T1 and T2 FLAIR after using on-scanner bias-field correction and T1’s FAST bias field correction. Figure 9 (panels A and B) shows the intensities for a typical subject, with similar improvement for T1 and T2 FLAIR evident. Figure 9.C shows the different inter-quartile-ranges for all 78 subjects, with the results further summarised in Table 2.

### 3.4. swMRI pipeline

The processing pipeline for this modality is shown in Figure 10.

Combining phase images across coils requires care due to anomalous phase transitions in regions of focal signal dropout for a given coil. Currently, all coil channels are saved separately to enable combination of phase images in post-processing. Each coil channel phase image is first high-pass filtered to remove low-frequency phase variations (including both coil phase profiles and field distortion from bulk shape). A combined complex image is generated as the sum of the complex data from each coil (unfiltered magnitude and filtered phase), and the final phase image is the phase of the summation of the second echo, since it has greater venous contrast. Careful inspection of a small number of subjects found no anomalous phase transitions from individual channels in the final combined image.

Venograms were calculated using an established reconstruction (Haacke et al., 2004), in which magnitude images are multiplied by a further filtering of the phase data to enhance the appearance of veins. The filtering consists of thresholding the phase image to set the phase to zero in voxels indicating diamagnetic susceptibility, and to take the phase to the fourth power in voxels indicating paramagnetic susceptibility. The chosen power represents a trade-off between venous-tissue contrast and noise in the phase data. This enhances image contrast around veins, which are the source of strongest paramagnetic susceptibility.

**Figure 7:**
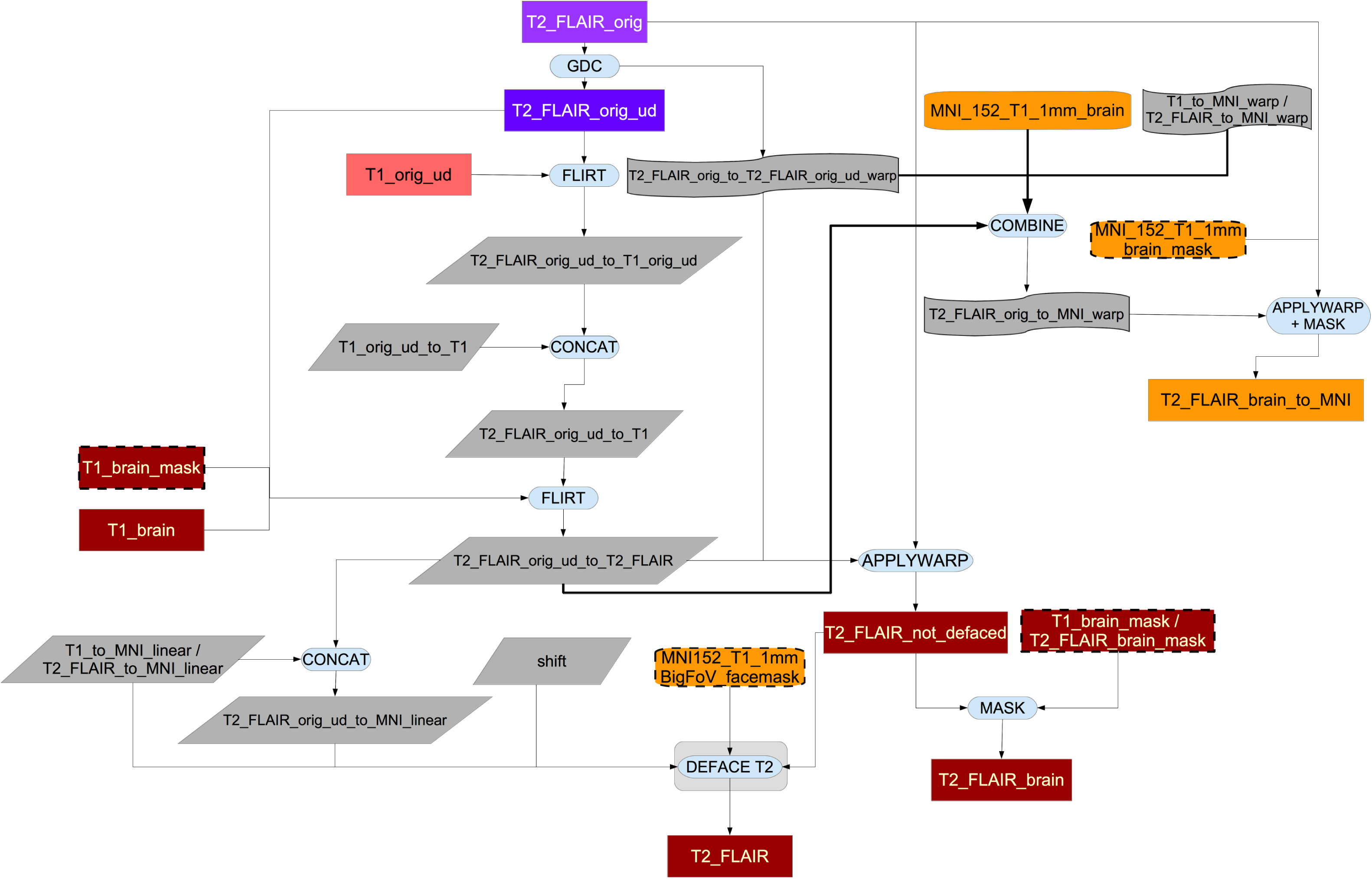
Flowchart for the T2 FLAIR processing pipeline.

**Figure 8:**
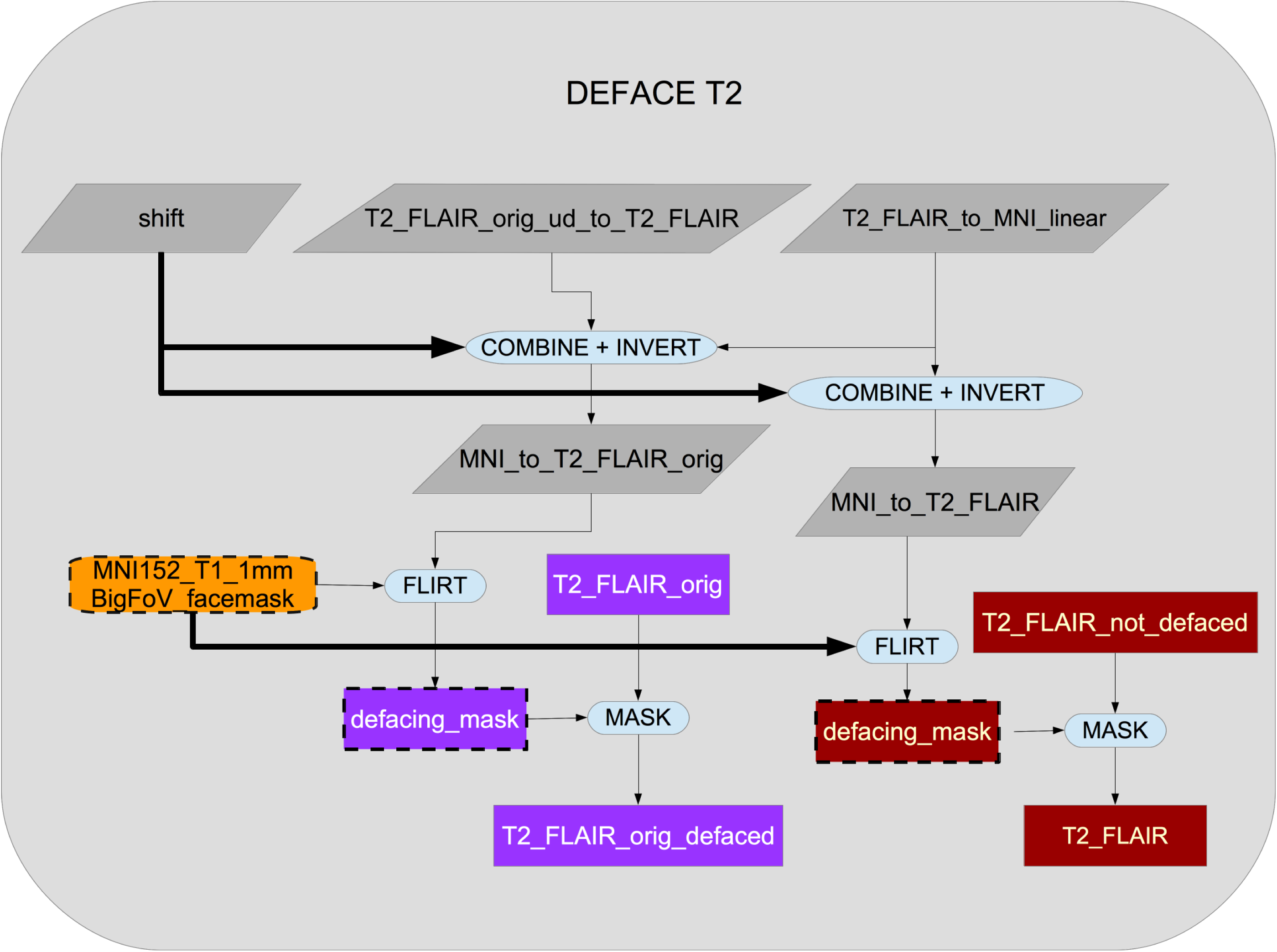
Flowchart for the defacing of the T2 FLAIR

**Figure 9:**
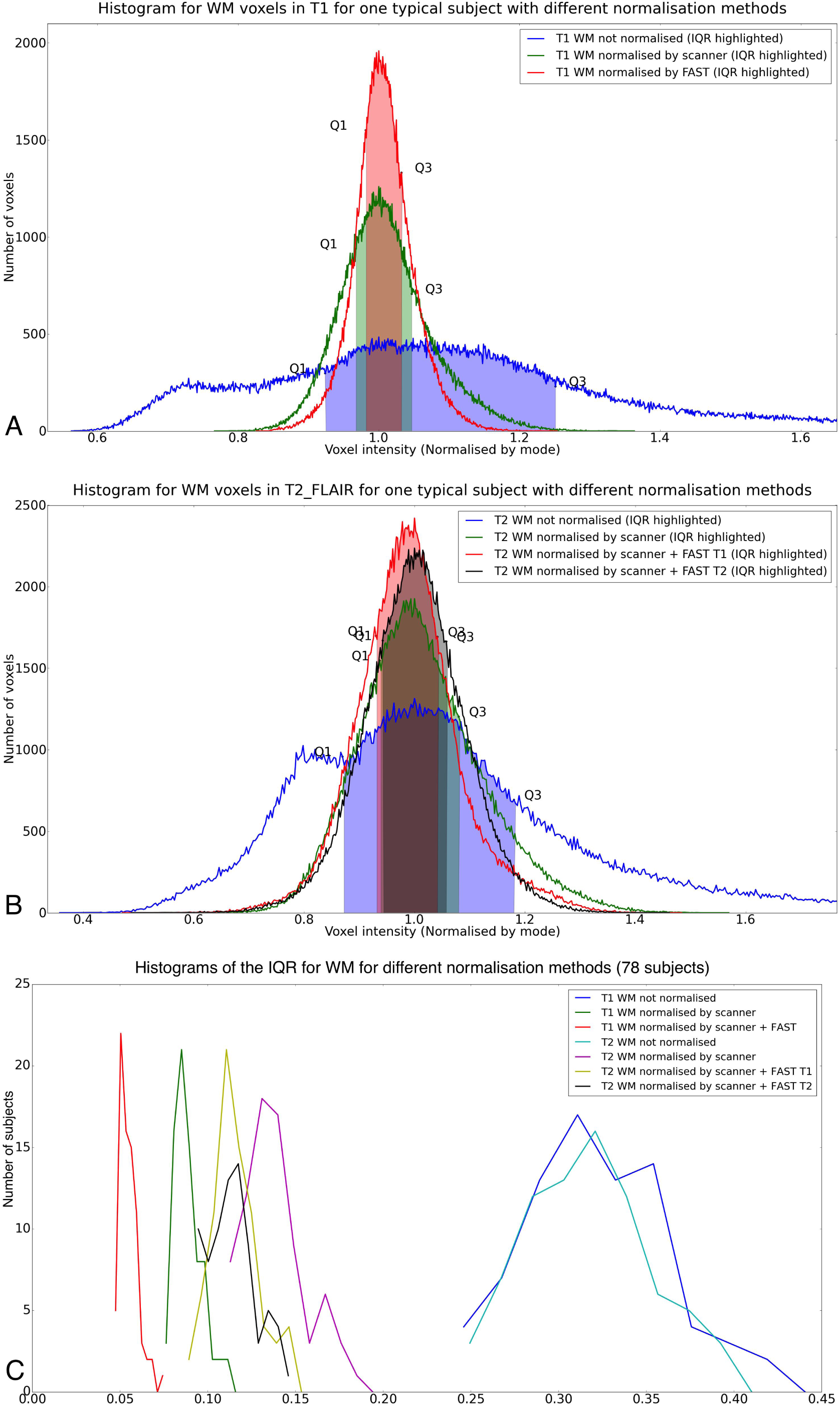
Histogram of WM intensities for a typical subject, showing that use of the bias field estimated from the T1 image can be applied to the T2 FLAIR image with similar improvement in IQR range to the one obtained in T1. Also, the histogram of this method is not very different to the one obtained by applying FAST directly to T2 FLAIR. *A:* T1. *B:* T2 FLAIR. *C:* Distribution of WM intensity Inter Quartile Ranges (IQRs) using different normalization methods in 78 subjects.

**Figure 10:**
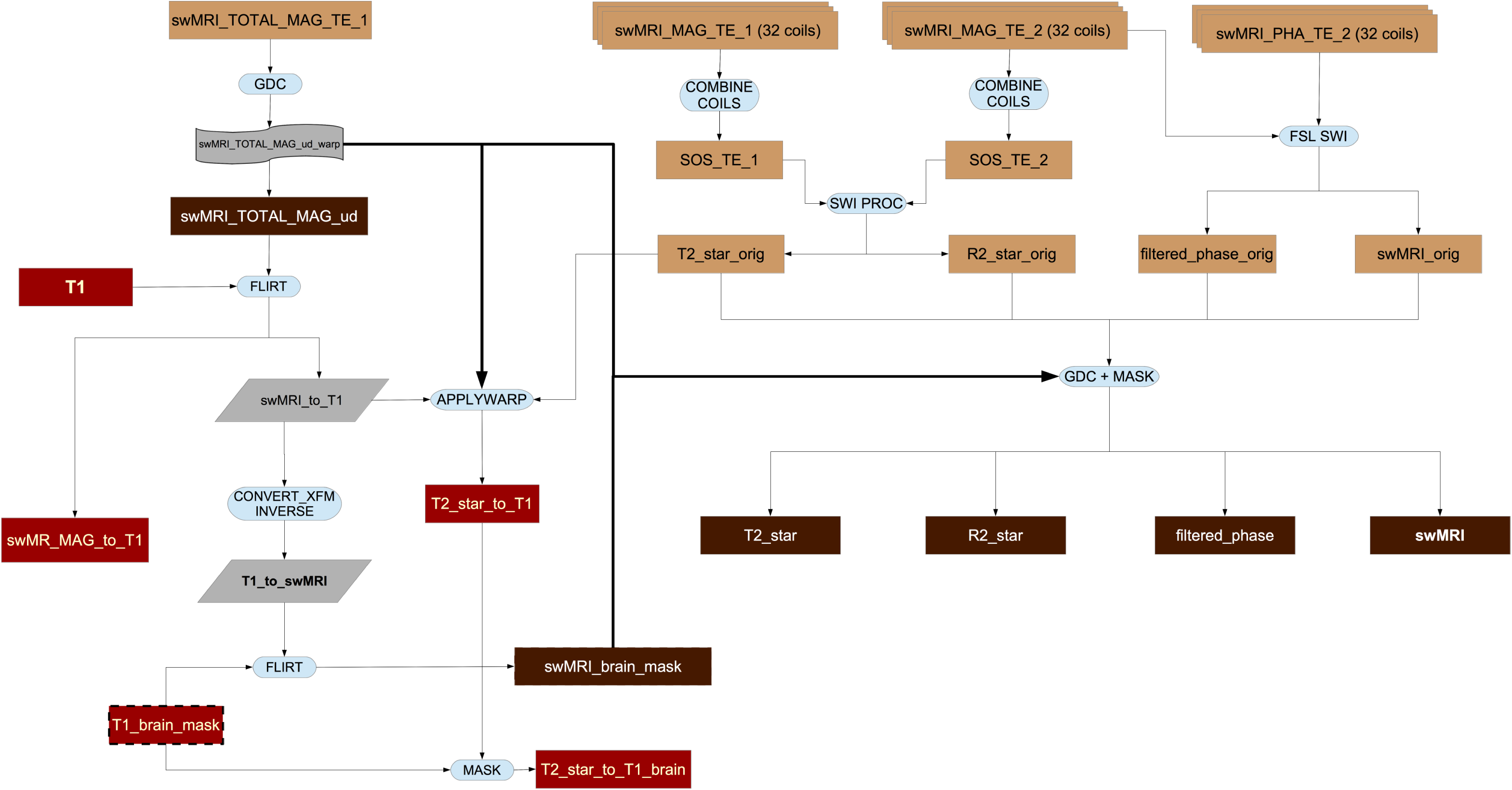
Flowchart for the swMRI processing pipeline.

**Table 2:** Descriptive statistics of WM intensity Inter Quartile Ranges (IQRs) using 3 normalisation methods (78 subjects).

| Normalisation method | IQR<br>Typical<br>Subject | Mean<br>IQR | Median<br>IQR | Std<br>IQR |
| --- | --- | --- | --- | --- |
| T1 not normalised | 0.3274 | 0.3232 | 0.3191 | 0.0418 |
| T1 normalised by scanner | 0.0797 | 0.0893 | 0.0872 | 0.0086 |
| T1 normalised by scanner + FAST | 0.0508 | 0.0550 | 0.0535 | 0.0058 |
| T2 FLAIR not normalised | 0.3106 | 0.3181 | 0.3163 | 0.0365 |
| T2 FLAIR normalised by scanner | 0.1429 | 0.1386 | 0.1356 | 0.0181 |
| T2 FLAIR normalised by scanner + FAST from T1 | 0.1116 | 0.1162 | 0.1138 | 0.0136 |
| T2 FLAIR normalised by scanner + FAST from T2 FLAIR | 0.1154 | 0.1144 | 0.1123 | 0.0137 |

Additionally, T2* values are calculated from the magnitude data. First, the inverse of T2*, termed R2*, is calculated from the magnitude images from the two echo times (TE1 and TE2). The log of the ratio of these two echo time images is calculated, and scaled by the echo time difference, to give the R2*.

T2* is calculated as the inverse of R2*. The T2* image is then spatially filtered to reduce noise (3x3x1 median filtering followed by limited dilation to fill small holes of missing data) and transformed into the space of the T1 (via linear registration of the bias-field-normalised first-echo magnitude image). The median (across ROI voxels) T2* value is then estimated as a separate IDP for each of the subcortical structure ROIs (left thalamus, right caudate, etc.) obtained from the T1.

### 3.5. Fieldmap generation pipeline

Fieldmap images reflect variations in the static magnetic field. These are needed to correct geometric distortions in the phase encoding direction in EPI images (Jezzard and Balaban, 1995).

In the fieldmap generation pipeline (see Figure 11) we estimate the fieldmaps from the b=0 images^19^ in the dMRI data using the opposing AP - PA phase-encoding acquisitions mentioned above. The reasons for doing so, rather than measuring the field using a dual echo-time gradient echo acquisition, are numerous. Firstly, “traditional” fieldmaps (based on dual echo-time gradient-echo images) are not free of problems (Andersson and Skare, 2010). The choice of echo-time difference represents a trade off between wanting a large echo-time difference to get a large effect to calculate the field from, and wanting a small echo-time difference which results in fewer phase-wraps that may be hard to unwrap. Furthermore, such a fieldmap will not automatically be in the same space (position) as the images one wants to correct. Therefore the fieldmap needs to be registered to those images and that is not a trivial task as they are differently distorted. Also, as mentioned in Smith et al. (2013), fieldmaps estimated with reversed encoding direction acquisitions produce an “equivalently good distortion correction accuracy” to that achieved with traditional fieldmaps, while being faster to acquire. This reduction in time, aside from making the acquisition less susceptible to within-scan head motion, saves acquisition time, which is a valuable resource in UK Biobank.

**Figure 11:**
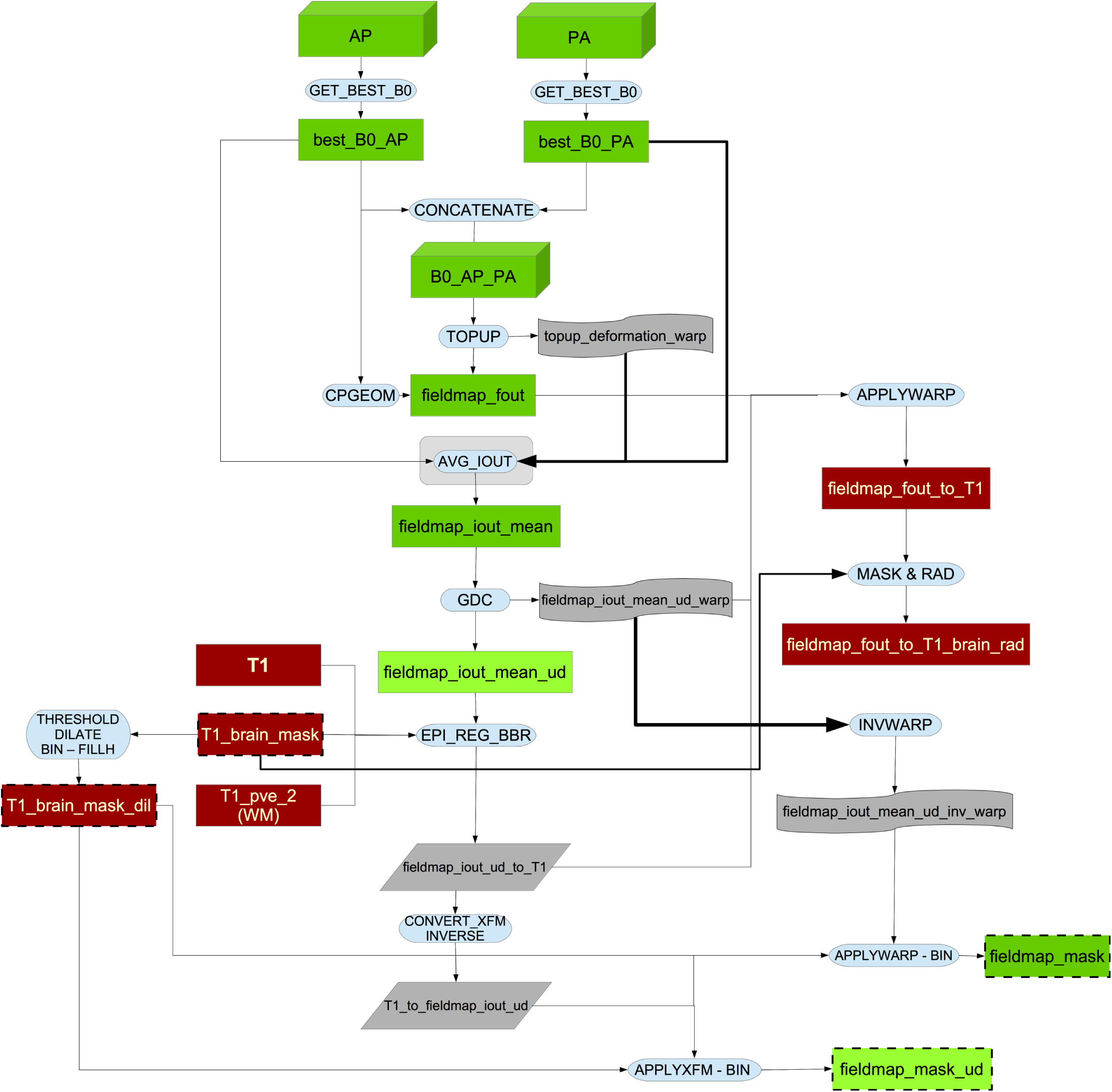
Flowchart for the fieldmap generation pipeline.

All b=0 dMRI images (interspersed with the high-b images) with opposite phase-encoding direction (AP and PA) are analysed to identify the most suitable pair of AP and PA images^20^. This is achieved by aligning all AP images to each other with a rigid-body registration (6 degrees of freedom) and then calculating the correlation of each b=0 image with all others. The volume that best correlates with all others is selected^21^ as the “target”, subject to the following additional criterion.

As the chosen b=0 image determines the space for later processing we want that space to be as close to the space of the first image with diffusion encoding (b>0) as possible. The movement estimation between the b=0 image (no diffusion weighting) images and the two shells with diffusion encoding is not carried out until the end of the eddy current correction, so if the fieldmap is far away from that first dMRI, the movement estimation process will be less optimal. Furthermore, dMRI images are acquired shortly after the fMRI images, so the case for selecting an early b=0 image is stronger^22^. Hence, we apply a selection bias towards the first b=0 image: if the first b=0 image has sufficient quality (correlation of 0.98 or higher to the other images) we would select it as the “best b=0 image” (the same selection bias is applied to PA).

This optimal AP/PA pair is then fed into topup^23^ (Andersson et al. (2003)) in order to estimate the fieldmap and associated dMRI EPI distortions.

The full generation method of the fieldmap magnitude is explained in section S5 of the supplementary material. GDC is applied to this magnitude image before registering it to the T1 (which has already been gradient-distortion corrected). The result is linearly aligned to the T1 (using boundary-based-registration [BBR] as the cost function as described in Greve and Fischl (2009)) for later use in unwarping the fMRI data. The resulting transformation is applied to the fieldmap and also inverted to get the dilated structural brain mask in fieldmap (dMRI) space. Finally, the fieldmap image in structural space is brain-masked and converted to radians per second for later use for fMRI unwarping.

### 3.6. dMRI pipeline

As can be seen in Figure 12, in the first step of this part of the pipeline the dMRI data (AP encoding direction) is corrected for eddy currents and head motion, and has outlier-slices (individual slices in the 4D data) corrected using the eddy tool (Andersson and Sotiropoulos, 2015, 2016; Andersson et al., 2016). This step requires knowing the “best” b=0 image in the AP direction as discussed above. The primary corrections carried out by eddy need to be done in-plane, and applying the GDC before eddy would move data out of plane. Therefore, GDC is applied after eddy to produce a more accurate correction, as explained in Glasser et al. (2013).

The output is fed into two complementary analyses, one based on tract-skeleton (TBSS) processing, and the other based on probabilistic tractography (bedpostx / probtrackx). Both analysis streams then report a range of dMRI-derived measures within different tract regions: A) measures derived from diffusion-tensor modelling, and B) measures derived from microstructural model fitting.

#### 3.6.1. Diffusion-Tensor-Image fitting and NODDI

The b=1000 shell (50 directions) is fed into the DTI fitting (Basser et al., 1994) tool DTIFIT, creating outputs such as fractional anisotropy, tensor mode and mean diffusivity.

**Figure 12:**
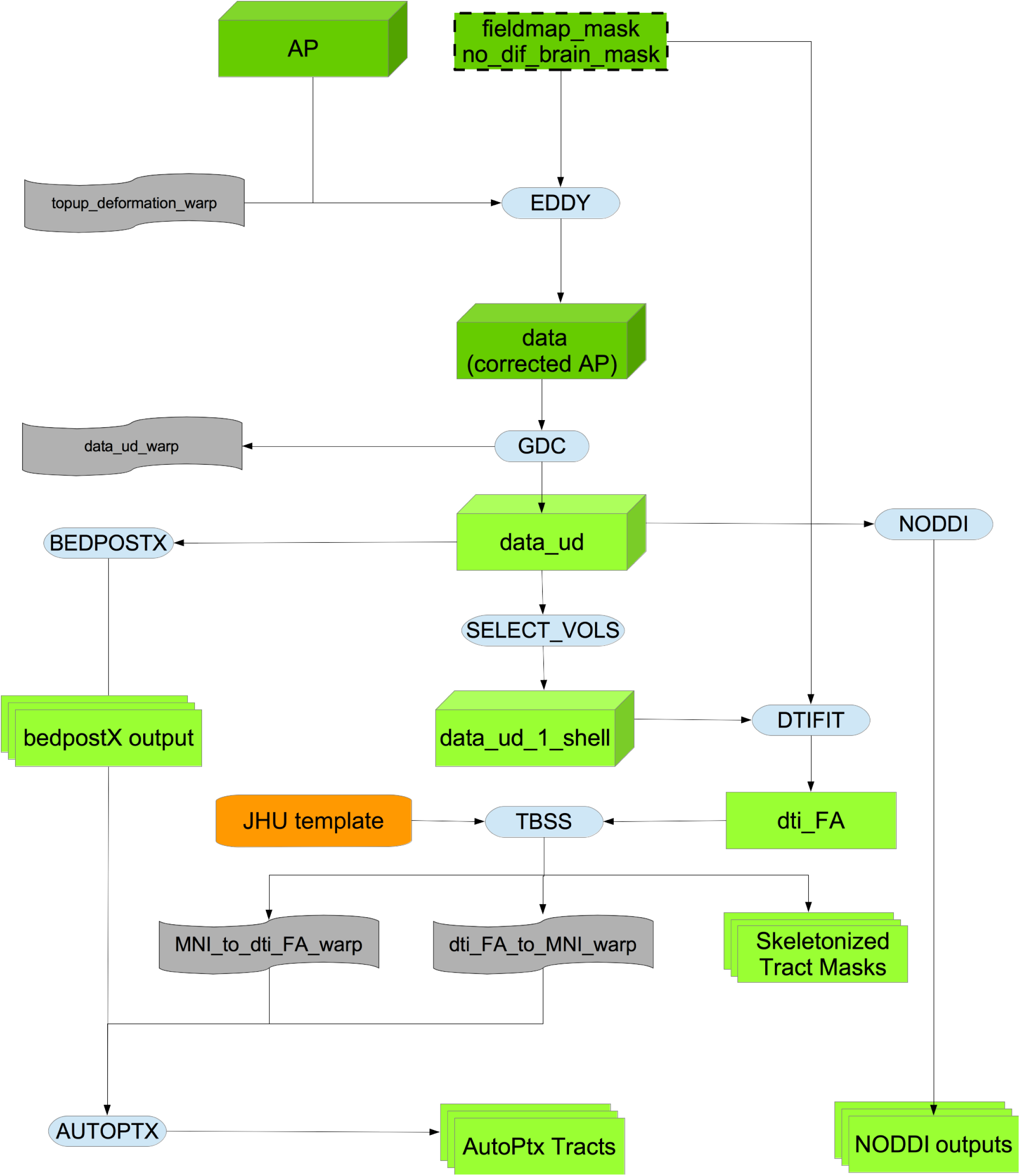
Flowchart for the dMRI processing pipeline.

In addition to the DTI fitting, the full two-shell dMRI data is fed into NODDI (Neurite Orientation Dispersion and Density Imaging) (Zhang et al., 2012) modelling, using the AMICO (Accelerated Microstructure Imaging via Convex Optimization) tool (Daducci et al., 2015). This aims to generate meaningful voxelwise microstructural parameters, including ICVF (intra-cellular volume fraction - an index of white matter neurite density), ISOVF (isotropic or free water volume fraction) and ODI (orientation dispersion index, a measure of within-voxel tract disorganisation).

#### 3.6.2. WM tract skeleton analysis

The DTI Fractional Anisotropy (FA) image is fed into TBSS (Smith et al., 2006), which aligns the FA image onto a standard-space white-matter skeleton, with alignment improved over the original TBSS skeleton-projection methodology through utilisation of a high-dimensional FNIRT-based warping (de Groot et al., 2013). This decision was re-validated along the lines of de Groot et al. (2013) after a thorough comparison of 14 different alignment methods applied to UK Biobank data (see Figure 13). The resulting standard-space warp is applied to all other DTI/NODDI output maps. For each of the DTI/NODDI maps, the skeletonised images are averaged within a set of 48 standard-space tract masks defined by the group of Susumu Mori at Johns Hopkins University Mori et al. (2005) and Wakana et al. (2007), (similar to the TBSS processing applied in the ENIGMA project Jahanshad et al. (2013)), to generate a set of dMRI IDPs.

#### 3.6.3. Probabilistic-tractography-based analysis

In addition to the TBSS analyses, the preprocessed dMRI data is also fed into a tractography-based analysis. This begins with within-voxel modelling of multi-fibre tract orientation structure via the bedpostx tool (Bayesian Estimation of Diffusion Parameters Obtained using Sampling Techniques)^24^, which implements a model-based spherical deconvolution and estimates up to 3 fibre orientations per voxel. This is followed by probabilistic tractography (with crossing fibre modelling) using probtrackx (Behrens et al., 2003, 2007; Jbabdi et al., 2012; Hernández et al., 2013). The bedpostx outputs are suitable for running tractography from any (voxel or region) seeding; the pipeline has already automatically mapped a set of 27 major tracts using standard-space start/stop ROI masks defined by AutoPtx de Groot et al. (2013). In order to reduce the amount of processing time for those tracts, AutoPtx was modified to reduce the number of seeds per voxel (the relationship between the number of seeds per voxel and processing time is linear) to 0.3 times the number of seeds specified in the original version of AutoPtx. Figure 14 illustrates the process we used to select this factor.

Although eddy and bedpostx outputs are in the space and resolution of the (GDC-unwarped) native diffusion data, the nonlinear transformation between this space and 1mm MNI standard space (as estimated by TBSS above) is used to create tractography results in 1mm standard space. For each tract, and for each DTI/NODDI output image type, we compute the weighted-mean value of the DTI/NODDI parameter within the tract (the weighting being determined by the tractography probabilistic output) to generate a further set of dMRI IDPs.

### 3.7. Resting fMRI pipeline

This section of the pipeline uses several outputs from the T1 pipeline (T1, brain-extracted T1, linear and non-linear warp of T1 to MNI152 space and binary mask of the white matter in T1 space). See Figure 15 for the flowchart.

Using this information and a previously calculated GDC warp for the rfMRI data, a configuration file is generated in a format suitable for use by Melodic (Beckmann and Smith, 2004). Melodic is a complete pipeline in itself which here performs EPI unwarping (utilising the fieldmaps as described above), GDC unwarping, motion correction using MC-FLIRT (Jenkinson et al., 2002), grand-mean intensity normalisation of the entire 4D dataset by a single multiplicative factor, and highpass temporal filtering (Gaussian-weighted least-squares straight line fitting, with sigma = 50.0s). To reduce interpolation artefacts, the EPI unwarping, GDC transformation, and motion correction are combined and applied simultaneously to the functional data. The combination of these warps can be seen in Figure 16.

**Figure 13:**
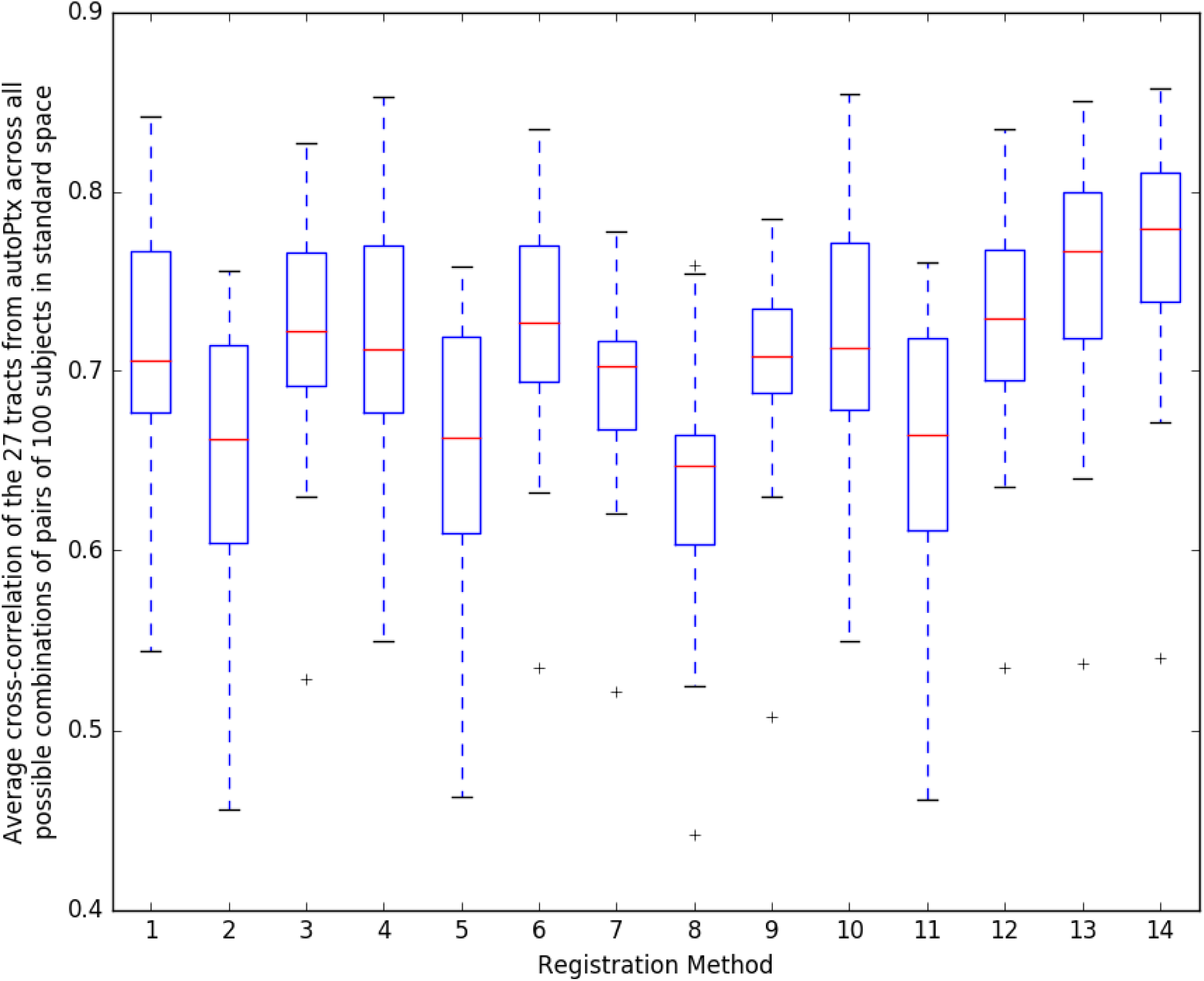
Comparison of 14 different alignment methods of FA to MNI space. We used the same methodology as de Groot et al. (2013). For each registration method, we used its estimated warp field in autoPtx to transform 27 automatically defined tracts into standard space; as discussed in de Groot et al. (2013), judging cross-subject alignment through similarity of tracts can be considered a test of alignment success that is reasonably independent of the images and cost functions used to drive the alignments.

*1:* FA linearly aligned to T1 + T1 non-linearly aligned to MNI.

*2:* FA linearly aligned to T1 + T1’s WM non-linearly aligned to MNI’s WM.

*3:* FA linearly aligned to T1 + T1’s GM non-linearly aligned to MNI’s GM.

*4:* Corrected b=0 linearly aligned (BBR) to T1 + T1 non-linearly aligned to MNI.

*5:* Corrected b=0 linearly aligned (BBR) to T1 + T1’s WM non-linearly aligned to MNI’s WM. *6:* Corrected b=0 linearly aligned (BBR) to T1 + T1’s GM non-linearly aligned to MNI’s GM. *7:* FA non-linearly aligned to T1 + T1 non-linearly aligned to MNI.

*8:* FA non-linearly aligned to T1 + T1’s WM non-linearly aligned to MNI’s WM. *9:* FA non-linearly aligned to T1 + T1’s GM non-linearly aligned to MNI’s GM. *10:* FA linearly aligned (BBR) to T1 + T1 non-linearly aligned to MNI.

*11:* FA linearly aligned (BBR) to T1 + T1’s WM non-linearly aligned to MNI’s WM.

*12:* FA linearly aligned (BBR) to T1 + T1’s GM non-linearly aligned to MNI’s GM.

*13:* FA non-linearly aligned to FA FMRIB58 atlas via an FA study-specific template (created by aligning all the FAs to FA FMRIB58 and then averaging).

*14:* FA non-linearly aligned to FA FMRIB58 atlas using high-dimensional FNIRT-based warping.

**Figure 14:**
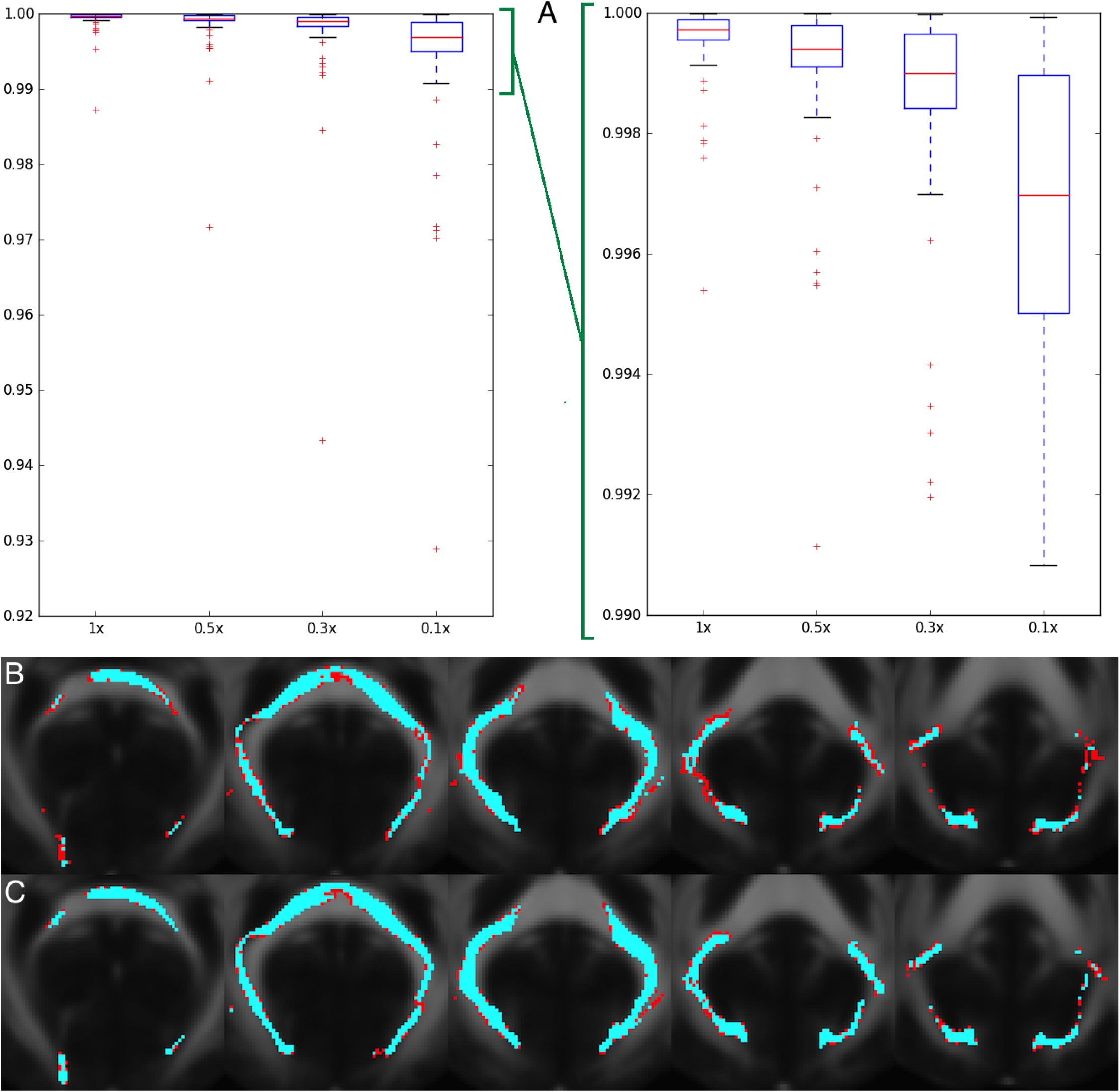
Degree of correlation and similarity of tracts after reducing the number of seeds per voxel in probtrackx. *A:* Average over 27 tracts and 5 subjects of the correlation between 2 different runs of probtrackx; this is reduced when we reduce the number of seeds per voxel. X axis is the reduction in the number of seeds with respect to the original AutoPtx configuration. Y axis is the average correlation of the pairs of tracts. When we reach 0.1x seeds per voxel, the median correlation drops below 0.999 for some tracts. Right plot is a zoom of left plot. *B:* Worst instance (in terms of correlation) of probtrackx in the forceps major tract with a factor of 0.1x seeds per voxel. The results remain robust even using a reduced number of seeds per voxel. Correlation between the maps for these two runs was **0.929**. Tracts were binarised for visualization using a threshold of 10% of the 99th percentile. The overlap of the 2 runs is shown in blue. The difference is shown in red. FMRIB58 FA atlas is shown in the background. *C:* Same tract (forceps major) from the same subject with a factor of 0.3x seeds per voxel. The results improve by increasing the number of seeds per voxel. Correlation between the maps for these two runs was **0.985**. FMRIB58 FA atlas is shown in the background. **The final decision was to use 0.3x.**

**Figure 15:**
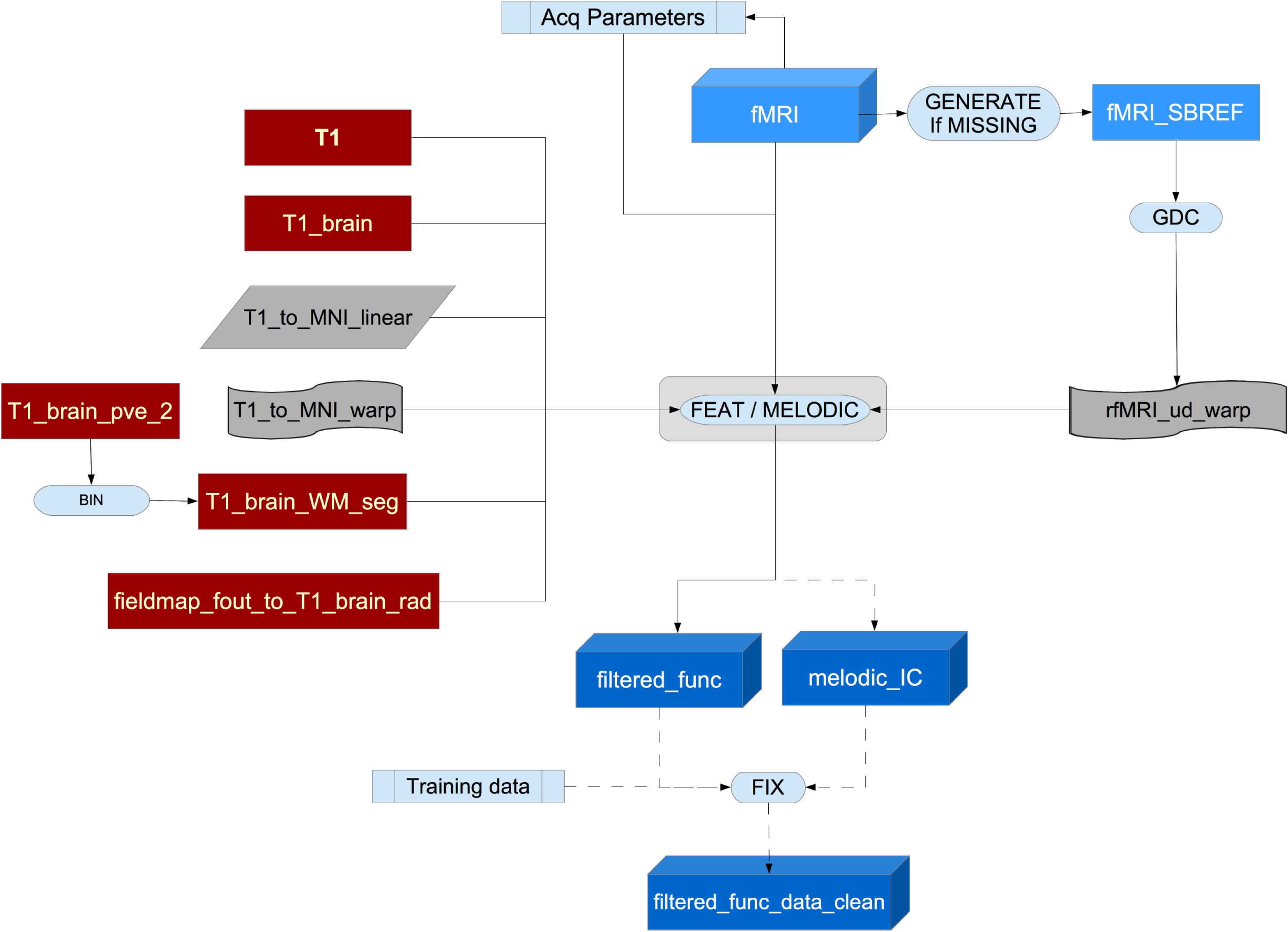
Flowchart for the fMRi processing pipeline.

Finally, structured artefacts are removed by ICA + FIX processing (independent component analysis followed by FMRIB’s ICA-based X-noiseifier (Beckmann and Smith, 2004; Salimi-Khorshidi et al., 2014; Griffanti et al., 2014). FIX was hand-trained on 40 Biobank rfMRI datasets following the methodology described in Griffanti et al. (2017), and leave-one-out testing showed (mean/median values across subjects) 99.1/100.0% classification accuracy for non-artefact components and 98.1/98.3% accuracy for artefact components. At this point no lowpass temporal or spatial filtering has been applied.

We evaluated the effect that FIX had on the relationship between 1/tSNR and amount of head motion (summarised to a single average value for each subject). Before FIX cleanup, the correlation was very high (r=0.75), indicating strong corruption of data in general by head motion, even after standard (geometric) head motion correction. However, after FIX cleanup, this correlation dropped dramatically (r=0.1, i.e., only 1% of variance explained by head motion), indicating high effectiveness of FIX in removing residual motion artefacts.

The EPI unwarping is a combined step that includes GDC and alignment to the T1, though the unwarped data is stored in native (unwarped) fMRI space (and the transform to T1 space stored separately). This T1 alignment is carried out by FLIRT, using BBR as the cost function. After the fMRI GDC unwarping, a final FLIRT realignment to T1 space is applied, to take into account any shifts resulting from the GDC unwarping. The previously described transform from T1 space to standard MNI space is utilised when fMRI data is needed in standard space.

To be able to identify RSNs (Resting-State Networks) in individual subjects, we first identify a set of RSNs which are common across the entire group. Therefore, group-average RSN analysis was carried out using 4100 datasets.

First, each subject’s preprocessed (as above) timeseries dataset was resampled into standard space, temporally demeaned and had variance normalisation applied according to Beckmann and Smith (2004). Group-PCA was then carried out by MIGP (MELODIC’s Incremental Group-PCA) from all subjects. This comprises the top 1200 weighted spatial eigenvectors from a group-averaged PCA (a very close approximation to concatenating all subjects’ timeseries and then applying PCA) Smith et al. (2014). The MIGP output was fed into ICA using FSL’s MELODIC tool Hyvärinen (1999); Beckmann and Smith (2004), applying spatial-ICA at two different dimensionalities (25 and 100^25^). The dimensionality determines the number of distinct ICA components; a higher number means that distinct regions within the spatial component maps are smaller. The group-ICA spatial maps are available online in the UK Biobank showcase^26^. The sets of ICA maps can be considered as “parcellations” of (cortical and sub-cortical) grey matter, though they lack some properties often assumed for parcellations - for example, ICA maps are not binary masks but contain a continuous range of values; they can overlap each other; and a given map can include multiple spatially separated peaks/regions. Any group-ICA components that are clearly identifiable as artefactual (i.e., not neuronally driven) are discarded for the network modelling described below. A text file is supplied with the publicly available group-ICA maps, listing the group-ICA components kept in the final network modelling. From the 25-dimensional group-ICA, 21 components were kept as non-artefactual, and from the 100-dimensional groupICA, 55 were kept.

**Figure 16:**
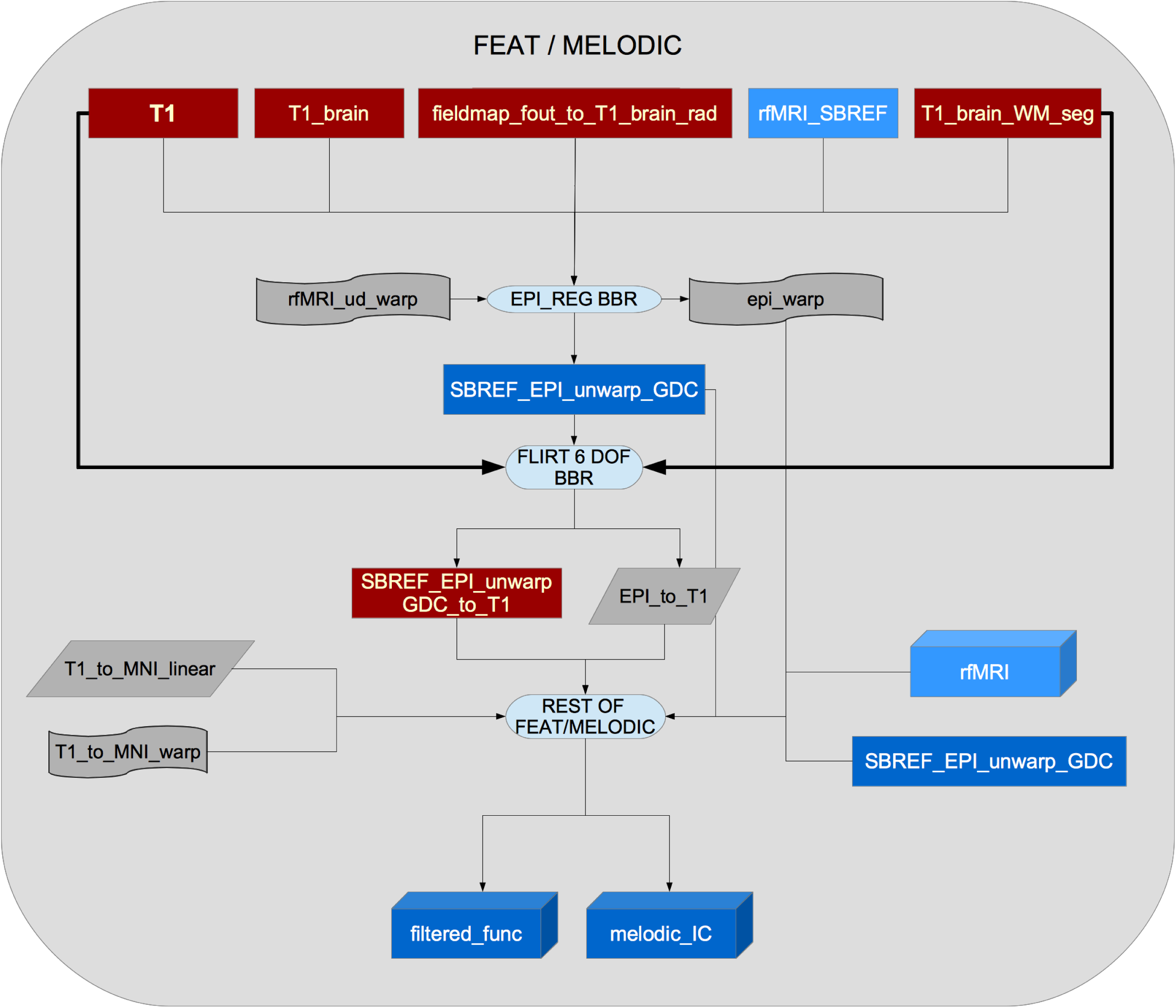
Flowchart for the FEAT part of the pipeline.

For a given parcellation (group-ICA decomposition of D components), the set of ICA spatial maps was mapped onto each subject’s rfMRI timeseries data to derive one representative timeseries per ICA component (for these purposes each ICA component is considered as a network “node”). For each subject, these D timeseries can then be used in network analyses, described below. This is the first stage in a dual-regression analysis Filippini et al. (2009).

The node timeseries are then used to estimate subject-specific network-matrices (also referred to as “netmats” or “parcellated connectomes”). For each subject, the D node-timeseries were fed into network modelling, after regressing the timeseries of the artefactual nodes out of all others, and then discarding them, leaving D_g_ nodes. This results in a *D*_g_ *× D*_g_ matrix of connectivity estimates. Network modelling was carried out using the FSLNets toolbox^27^. Network modelling is applied in two ways:

- Using full normalised temporal correlation between every node time series and every other. This is a common approach and is very simple, but it has various practical and interpretational disadvantages including an inability to differentiate between directly connected nodes and nodes that are only connected via an intermediate node (Smith, 2012), as well as being more corrupted (than partial correlation) by residual shared/global artefacts.
- Using partial temporal correlation between nodes’ timeseries. This aims to estimate direct connection strengths better than achieved by full correlation. To slightly improve the estimates of partial correlation coefficients, L2 regularization is applied (setting rho=0.5 in the Ridge Regression netmats option in FSLNets).

Netmat values were transformed from Pearson correlation scores (r-values) into z-statistics, including empirical correction for temporal autocorrelation. Group-average netmats are also available online.

As the matrices are symmetric, only values above the diagonal are kept, and unwrapped (taken column-wise) into a single row of D_g_ x (D_g_-1) / 2 values per subject. This results in one “compound” IDP (containing all network matrix values for a given subject) for each original dimensionality (D=25 and 100) and for each network matrix estimation method (full correlation and partial correlation).

### 3.8. Task fMRI pipeline

The same preprocessing and registration was applied as for the rfMRI described above, except that spatial smoothing (using a Gaussian kernel of FWHM 5mm) was applied before the intensity normalisation, and no ICA + FIX artefact removal was performed, both decisions being largely driven by the shorter timeseries in the tfMRI (than the rfMRI) and because of the greater general reliance in tfMRI analysis on voxelwise timeseries modelling (as opposed to multivariate spatiotemporal analyses common in resting-state fMRI).

Pre-processing and task-induced activation modelling was carried out using FEAT (FMRI Expert Analysis Tool); time-series statistical analysis was carried out using FILM with local autocorrelation correction Woolrich et al. (2001). The timings of the blocks of the two task conditions (shapes and faces) are defined in 2 text files. 5 activation contrasts were defined (Shapes, Faces, Shapes+Faces, Shapes-Faces, Faces-Shapes), and an F-contrast also applied across Shapes and Faces.

The 3 contrasts of most interest are: 1 (Shapes), 2 (Faces) and 5 (Faces-Shapes), with the last of those being of particular interest with respect to amygdala activation. Group-average activation maps were derived from analysis across all subjects, and used to define ROIs for generating tfMRI IDPs. Four ROIs were derived; 1 (Shapes group-level fixed-effect z-statistic, thresholded at Z>120^28^); 2 (Faces group-level fixed-effect z-statistic, thresholded at Z>120); 5 (Faces-Shapes group-level fixed-effect z-statistic, thresholded at Z>120); 5a (Faces-Shapes group-level fixed-effect z-statistic, thresholded at Z>120, and further masked by an amygdala-specific mask). The group-average activation maps and ROIs are available online in the UK Biobank showcase mentioned above.

The Featquery tool was used to extract summary statistics for these 4 contrast/mask combinations, for both activation effect size (expressed as a % signal change relative to the overall-image-mean baseline level) and statistical effect size (z-statistic), with each of these summarised across the relevant ROI in two ways - median across ROI voxels and 90th percentile across ROI voxels.

Display of the task video and logging of participant responses is carried out by ePrime software, which provides several response log files from each subject. These are not used in the above analyses (as the timings of the task blocks are fixed and already known, and the correctness of subject responses are not used in the above analyses), but are available in the UK Biobank database.

## 4. Automated Quality Control

We have developed an automated Quality Control tool to identify images with problems either in their acquisition or in later processing steps. This uses machine learning methods with supervised learning. We first categorise the different problems that we may find in the images and manually classify many datasets accordingly. We then generate a set of QC features aiming to characterise those images. Finally, we feed that information into a supervised learning classifier. With this approach we aim to have a classifier capable of detecting problematic images with acceptable accuracy. To date we have concentrated just on automated QC for the T1 images, as the entire processing pipeline depends on having a usable T1.

### 4.1. QC categories

Table 3 shows the categories for QC that were developed to classify issues that may be found in T1 images. This list was compiled as new issues were identified (through manual inspection) in the data, and is not intended to cover every possible problem in any MR image, but rather to cover all of the problems that we have observed to date in the Biobank data set. For this study, we define a “problem” as any issue that makes subsequent processing steps impossible or unreliable, while an “imperfection” is an issue which must be noted, but does not necessarily impede further processing. Some issues (e.g., bad head motion) appear in both categories (“problem” and the less serious “imperfection”), where the distinction is to be made on the basis of the seriousness of the image artefacts. Table S2 in the supplementary material shows a similar table for all modalities. Figure S9 in the supplementary material shows the difference in the IDPs between “problematic”, “imperfect” and “good” datasets). Figure S12 in the supplementary material shows some examples of some datasets from these categories.

### 4.2. QC features

We developed a set of 190 features (for use in the trained machine learning automated classifier) to find issues in T1 structural images. A more detailed description can be found in Section S4 of the supplementary material. The features that were developed are focused on characterizing:

- Discrepancy between the T1 (after alignment into MNI standard space) and the MNI template.
- Signal-to-noise ratio.
- Total brain volume and segmented tissue volumes.
- Subcortical structures, volumes.
- Asymmetry between the subcortical structures.
- Global brain asymmetry.
- Normalised intensity of the subcortical structures.
- Volume of grey matter (Using a different brain extraction method) outside the brain mask.

**Table 3:** Classification of problems / imperfections for T1

| Problem code | Imperfection code |
| --- | --- |
| 0 = No problem<br>1 = Multiple / Unknown problems<br>2 = Missing or incomplete modality<br>3 = Bad FOV<br>4 = Bad registration: bad head motion<br>5 = Bad registration: structurally atypical<br>6 = Bad registration: unknown reason | 1 = Multiple / Unknown imperfections<br>2 = Bad head movement<br>3 = Movement-related ringing / blurring<br>4 = Bias field / contrast problem<br>5 = Structurally atypical |

**Incomplete** = The expected number of images or any of the dimensions (including the temporal dimension) is incorrect.
**Structurally atypical:** The subject is anatomically atypical. This may be due to a pathology although there was no clinical assessment to confirm this. The problem may be so severe that registration is strongly affected (and thus, the image is deemed unusable for the automated processing pipeline).

- Amount of segmented tissue in the border of the brain mask.
- “Volume” of the edges (derived using Canny filter and excluding certain borders) of each segmented tissue.
- Comparison with different brain extraction tools.
- Intensity in the internal and external border of the brain mask.
- Volume of holes in grey and white matter.
- Magnitude of warp field from the non-linear registration to the MNI template.
- Volume of White Matter Hyperintensities.
- Distance (per lobe) between the border of the brain mask and the border of the MNI template.

### 4.3. Automated QC tool

We used the Weka machine learning toolbox (Hall et al., 2009), with 3 separate classifiers’ outputs fused together. The ensemble classifier used for the fusion was a voting system that combines the a posteriori probabilities of the different classifiers using the ‘Minimum Probability’ combination rule (Kuncheva, 2004; Kittler et al., 1998). This rule was chosen^29^ among 5 different options based on our goal of reducing the FNR (False Negative Rate: rate of missing “problems”) as much as possible without increasing the FPR (False Positive Rate: rate of incorrectly flagging “good” cases as “problems”) to impractical levels^30^. The three combined classifiers were:

- Bayes Network classifier (Bouckaert, 2008): This classifier first finds the Bayesian network (as a directed acyclic causal graph) that best fits the data by using the training dataset to create the network structure (running conditional independence tests to find the causal structure of the network, using local score metrics on quality of nodes/edges, and performing global score metrics by estimating classification accuracy) and then assigning probabilities to each node using the distribution of the same training dataset; the inference is later performed by using the maximum a posteriori decision rule for the node of interest (i.e., the variable being predicted) for a new instance (a vector of features).
- Naive Bayes classifier (John and Langley, 1995): This classifier is a simpler version of the above one in the sense that it assumes strong (naive) independence between the features (in the sense that there is no correlation between the different features) and therefore does not need to generate a graph structure. The maximum a posteriori decision rule for the parent node is also used here for the inference.
- MetaCost classifier (Domingos, 1999): MetaCost is a meta-classifier that allows the use of different costs for penalising (e.g.) FPR vs FNR. for a certain base classifier. In our case, as the primary goal was to try to minimise the False Negatives (subjects that are actually unusable but were classified as usable) we created a cost matrix that penalised greatly this kind of misclassification. This cost matrix was calculated using a grid search of different parameters and can be seen in Table S1 of the supplementary material. The base classifier was another Bayes Network classifier.

Combining classifiers usually tries to compensate for possible overfitting (or lack of fitting) of individual classifiers. The reason for choosing this combination of algorithms was that they allowed us to weigh the importance of type I vs type II errors, besides exhibiting a satisfactory accuracy in earlier stages of the classification (again, with a reduced version of the current training set).

**Table 4:** UK Biobank usable modalities in acquisition order. Total subjects: 10,129.

| Modality | Available | Usable |
| --- | --- | --- |
| T1 | 10,102 | 9,933 (98.06%) |
| rfMRI | 10,034 | 9,558 (94.36%) |
| tfMRI | 9,620 | 9,182 (90.65%) |
| T2 FLAIR | 9,756 | 9,045 (89.30%) |
| dMRI | 9,566 | 8,839 (87.26%) |
| swMRI | 9,396 | 9,153 (90.36%) |
| All | 8,996 | 8,211 (81.06%) |

Very early improvements in the dMRI and T2 FLAIR protocols were found to be valuable, resulting in large enough data improvements to outweigh the priority of keeping the protocol fixed (and taking into account the relatively small numbers of datasets affected). This change was made at the beginning of protocol.

## 5. Results

### 5.1. UK Biobank as a resource

So far, the pipeline has been applied to 10,129 subjects. In November 2015, there was a public release of the imaging datasets and IDPs from the first 5847 subjects (Miller et al., 2016); a new set of 4282 subjects has recently been processed and publicly released in February 2017 (Biobank, 2017). The number of datasets from every modality that were deemed suitable for processing are listed in Table 4.

Expanding on the analyses made in Miller et al. (2016), we have run ∼8 million univariate correlations between IDPs and non-brain-imaging variables, to illustrate the statistical power of this resource. This analysis (see Figure 17.A) not only shows results largely compatible with the ones presented in Miller et al. (2016), but also that correlations with a P value smaller than 10^−100^ are possible due to the large number of subjects. Figure 17.B demonstrates more directly that the new set of 4282 subjects show very similar characteristics to the first set of 5847 subjects.

Figure 18.A shows a very simple example of the biological relevance of IDPs. In this case, the volume of White Matter Hyperintensities (WMHs) can be used to explore the relationship between age and WMHs. Figure 18.B shows an example of the WMH segmentation using BIANCA. This metric also correlates with known health biomarkers, such as systolic and diastolic blood pressure. The positive correlation between total WMH volume and systolic blood pressure (after correcting for the usual confounds^31^) has a Bonferroni-corrected significance of *P* < 10^−20^, *r* = 0.13, while the correlation with diastolic blood pressure has a Bonferroni-corrected significance of *P* < 10^−15^, *r* = 0.11. These significant associations are consistent with the literature (Liao et al., 1996; Gunstad et al., 2005).

**Figure 17:**
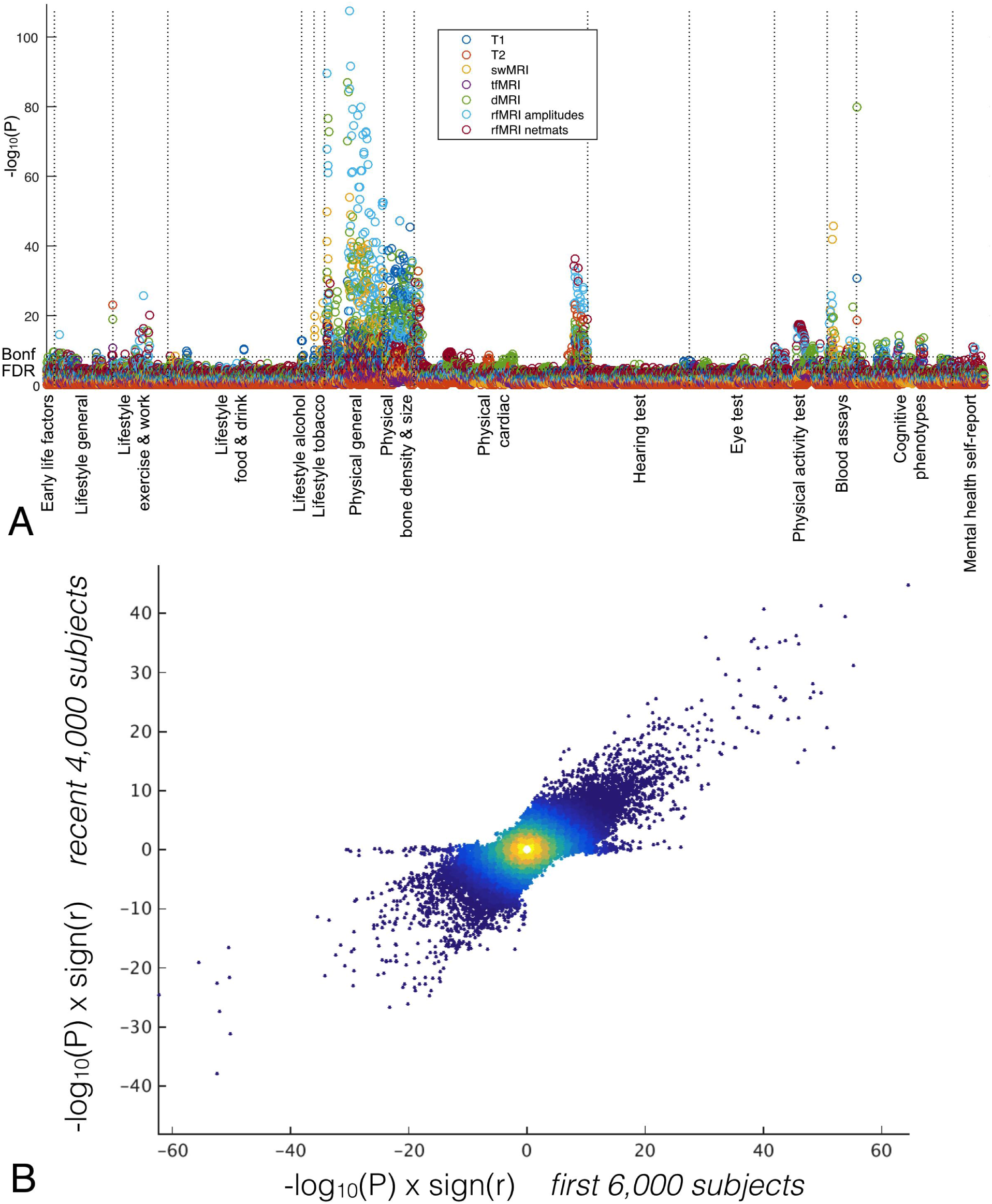
*A:* Manhattan plot summarising the significance of 8 million univariate association tests between IDPs and non-brain-imaging variables in the UK Biobank database from 10,000 subjects. For each non-imaging variable (i.e., each column in the plot), only the strongest association is plotted for each class of IDP, for clarity. Plotted p-values are not corrected for multiple comparisons, but the thresholds for both false-discovery-rate and Bonferroni correction are shown as dotted lines. *B:* High reproducibility (r=0.62) of these associations in the original vs. new groups of subjects; each point is a given IDP - non-brain-variable pairing. The small number of points along the y=0 axis relate to a non-imaging measure which (as a result of investigating these points in this plot) was found to be badly drifting over time).

**Figure 18:**
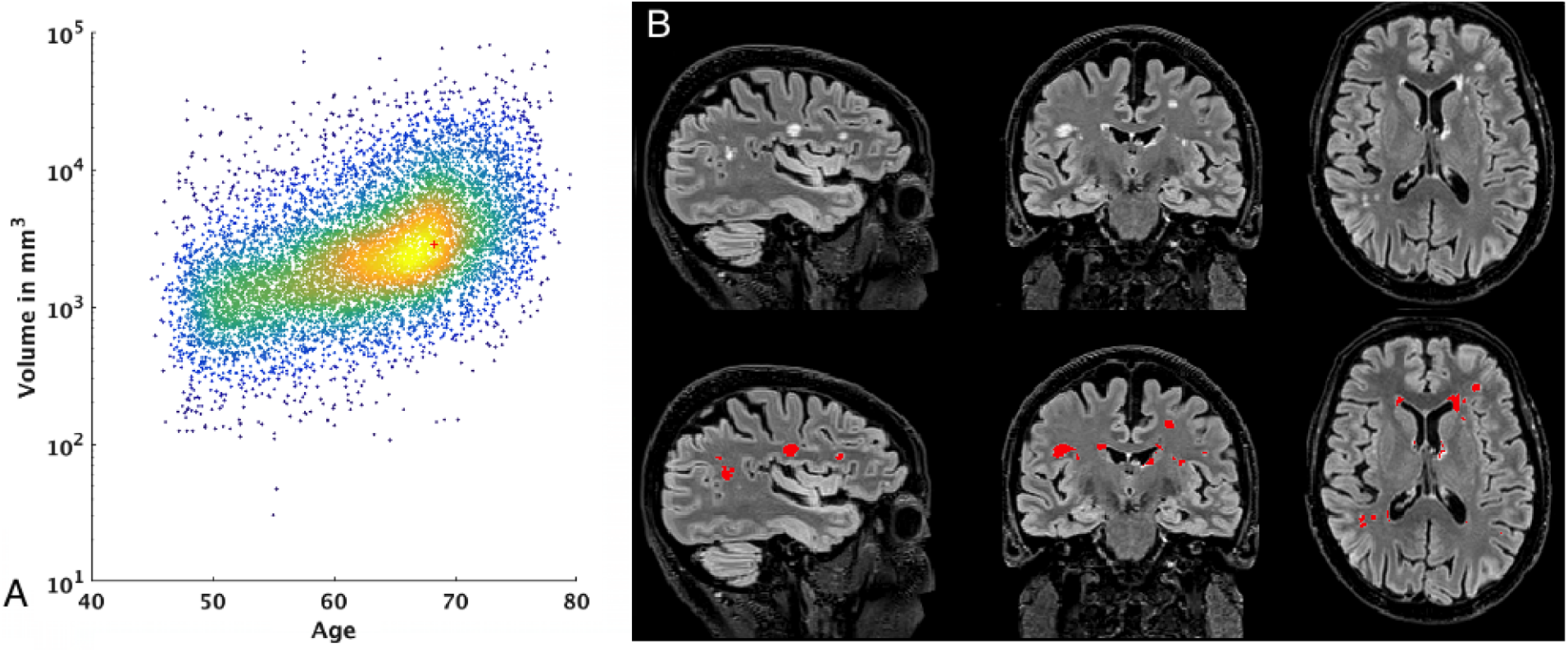
*A:* Relationship between age and total volume of white matter hyperintensities. Red cross shows the subject on the right. *B:* WMH segmentation using BIANCA on one example dataset Age: 68.5 years. Total WMH volume: 5049 *mm*^3^.

**Table 5:** Automated QC tool performance for T1 images

|  | Positive | Negative |  |
| --- | --- | --- | --- |
| **Classified Positive** | TP: 90 | FP: 747 | P. Precision: 11% |
| **Classified Negative** | FN: 8 | TN: 5002 | N. Precision: 99.8% |
|  | Sensitivity: 92% | Specificity: 87 % |  |

**Positive** = Unusable datasets. **Negative** = Usable datasets.
**Datasets flagged for manual review:** 837 / 5847 (14.31 %)
**Unusable datasets missed:** 8 / 5847 (0.13 %)

### 5.2. Performance of the automated QC tool

In order to validate our QC system, image quality for 5847 subjects was manually assessed. Problems were identified in 98 of these subjects. Table 5 shows the performance of the classifier in a stratified^32^ 10-fold cross validation.

The majority of problems found in the processing are related to non-linear registration, which likely therefore also affect the brain extraction. The 98 subjects were deemed unusable due to irreconcilable problems with non-linear registration, in many cases caused by poor data quality (for example, where the T1 is corrupted by bad subject head motion, as can be seen in Figure S12, panel 4 in the supplementary material). The QC system we have developed aims to detect problems like this.

The number of false negatives (that is, subjects that are actually unusable but were classified as usable), is low (8 subjects, i.e. 0.13%), though in future work we will aim to reduce this even further. Choosing parameters/thresholds to drive this number of false negatives so low comes at the cost of a much higher false positive rate (747 / 5749 = 13%). Nevertheless, these results guarantee a considerable reduction in the number of subjects in future data sets that will then require manual QC assessment (i.e., manually checking 837 subjects instead of 5847).

This automated QC tool was used on the second release of 4282 subjects, significantly reducing the manual checking to approximately 750 subjects, resulting in the detection of 71 unusable datasets.

## 6. Discussion and future work

The UK Biobank brain imaging data and IDPs can be used to create models that describe the population, and in combination with the healthcare outcomes (which will feed into UK Biobank over coming years), be able to identify risk factors and biomarkers for early diagnosis of many diseases. In this paper we provided the first detailed description of the core processing pipeline being developed to process the raw imaging data and generate the IDPs. We also described several investigations necessary to feed into decisions made during the development of the acquisition protocol and the analysis pipeline.

The huge number of subjects makes the need for a completely automated tool evident, including the detection of inadequate datasets or failures in processing. For this reason, the development of an adequate automated QC tool is an absolute necessity, and we have presented here an approach for automated QC of T1 data.

The study of possible drifts in imaging data over time will be important, and will be the subject of a future paper. We will investigate the possible effects of minor scanner software upgrades and minor protocol changes, as well as inter-site imaging consistency. We will also investigate the extent to which simple adjustment for such drifts may be effective.

A long-term future challenges will be data analysis and population modelling. The application of new unsupervised learning methods for finding new data-driven IDPs or QC metrics is potentially interesting (Duff et al., 2015). This could also evolve to consider health outcomes as noisy / weak labels (Fréenay and Verleysen, 2014) in such a way that the learning process considers their uncertainty and reliability.

Future versions of the processing pipeline will incorporate new functionalities from other sources into the pipeline (such as the HCP, Glasser et al. (2013)). This will include surface generation via Freesurfer (Dale et al., 1999), adding morphometric measures such as cortical thickness or cortical surface area, mapping resting fMRI and dMRI onto the cortical surface, and using Multimodal Surface Matching (Robinson et al., 2014) for surface-based registration. As the resolution, quality and quantity (e.g., number of fMRI timepoints) in UK Biobank data cannot match that in the HCP (with its 4 hours of scanning per subject) each of these steps will involve careful evaluation and potential reworking in the context of our data. For example, projection of the lower-resolution fMRI data onto the cortical surface vertices, including the HCP approach of removing “noise” voxels, will need careful optimisation.

Another new path to explore would be evaluating how using different modalities for some processing steps may improve some results. These operations may include: using T1 + FLAIR versus T1 in Freesurfer; using T1 + FLAIR versus T1 in FAST; using T1 + FLAIR + dMRI versus T1 + FLAIR in BIANCA.

The development of the QC tool allowed us to find problematic steps in the processing pipeline (i.e., non-linear registration to the MNI template). Even though the number of subjects with this problem is not large (98/5847 = 1.7%, compared to the 2-13% failure rate for the acquisitions), and we can detect them fairly consistently, improving this step will be a future line of work. Possible improvements may involve a combination of the following:

- Generating a study-specific T1 template and registering subject images to it instead of to the MNI template.
- Improving brain extraction and using the resulting brain mask to guide registration. Possibilities include running FNIRT with much lower degrees of freedom to initialise brain extraction, and/or developing a classifier that works directly on each voxel to classify them as brain/non-brain. See the Appendix for more details on the brain extraction method we have used to date.
- Combining T1+dMRI (and possibly more modalities) in the registration in a more integrated manner; for example, taking advantage of the richer signal within white matter available from the diffusion imaging.

A possible way to improve QC accuracy would be in the development of better features to drive the classification.

These new features may be created by finding new heuristics that better describe the problems or by using unsupervised machine learning techniques to model the data, e.g., using ICA-like techniques (Hyvärinen, 1999; Beckmann and Smith, 2004).

In addition, we need the development of metrics to describe the discrepancy between the T1 image (for a given subject) and each of the other modalities (for that same subject), after linear alignment of the other modalities to the T1. Therefore, we are planning to extend the automated QC system to reliably find problems in the acquisition and processing steps for all modalities (not just T1). Again, ICA-like techniques may be useful here, as well as methods based on the Hidden Markov Model (Baker et al., 2014; Vidaurre et al., 2016) to interrogate the network dynamics.

Having a fixed acquisition protocol and a core processing pipeline in UK Biobank is a key feature of this project (Gronenschild et al., 2012; Glatard et al., 2015). Providing a core pipeline that generates both processed data (e.g., with images artefact-cleaned and aligned across modalities and subjects) as well as IDPs will hopefully be valuable to both imaging researchers and non-imaging experts (healthcare researchers, epidemiologists, etc.). Having said that, the intention is not to discourage other imaging researchers from developing their own image processing and IDP generation approaches and software; indeed, UK Biobank encourages all researchers to feed quantitative research outputs back into the central database. Considering the invaluable connection of the outputs from processing the brain imaging data with the rest of UK Biobank resources and NHS records, the possibilities for future research are massive.

## Acknowledgements

UK Biobank brain imaging and FAA are funded by the UK Medical Research Council and the Wellcome Trust. LG is funded by the NIHR Oxford Biomedical Research Centre and the Monument Trust Discovery Award from Parkinson’s UK. GD is supported by the UK Medical Research Council (MRC) MR/K006673/1. EV is supported by the Engineering and Physical Sciences Research Council (EPSRC) D4TD4D12/HM12.01. The authors gratefully acknowledge funding from the Wellcome Trust UK Strategic Award [098369/Z/12/Z]. The Wellcome Centre for Integrative Neuroimaging is supported by core funding from the Wellcome Trust (203139/Z/16/Z). PMM thanks the Edmond J. Safra Foundation, Lily Safra, NIHR and the Imperial College Healthcare Trust BRC for personal support. Additional input on methods, processing pipeline and informatics: Matteo Bastiani, Eugene Duff, Sean Fitzgibbon, David Flitney, Steve Garratt, John Miller, Duncan Mortimer, Jonathan Price, Gholamreza Salimi-Khorshidi, Anderson Winkler and Alan Young. Additional input on machine learning by Pradeep Reddy Raamana. Additional input on data visualization by Mike Lawrence.

## 7. BIBLIOGRAPHY

## Footnotes

1 The raw and processed imaging data, IDPs and non-imaging measures in UK Biobank are available to researchers worldwide following a data access application procedure.

2 http://www.fmrib.ox.ac.uk/datasets/ukbiobank/nnpaper/IDPinfo.txt.

3 A comprehensive list of possible artefacts and proposed qualitative quality recommendations can be found in CBS (2016).

4 http://preprocessed-connectomes-project.org/quality-assessment-protocol/

5 https://amsportal.ukbiobank.ac.uk/sitepages/sign%20in.aspx.

6 Manual QC was possible because at that stage, we were dealing with the initial phase (6% of final dataset size)

7 GPU processing is used for the FSL eddy and bedpostx tools, with further tools being ported to GPU in the future

8 https://git.fmrib.ox.ac.uk/falmagro/UK_biobank_pipeline_v_1

9 http://www.fmrib.ox.ac.uk/ukbiobank

10 See section 3.2.

11 33 out of 5822.

12 https://github.com/Washington-University/Pipelines.

13 More information about this file can be found in: <u>https://github.com/Washington-University/Pipelines/wiki/FAQ</u>

14 Default cost function: Correlation ratio.

15 http://www.bic.mni.mcgill.ca/ServicesAtlases/ICBM152NLin6.

16 Configuration file for optimal registration is distributed with the pipeline code.

17 Structural Image Evaluation using Normalisation of Atrophy: Cross-sectional.

18 The ROIs are defined by combination of parcellations from several atlases: Harvard-Oxford cortical and subcortical atlases, and Diedrichsen cerebellar atlas

19 b=0 images (also called b0) have a b-value below 50.

20 Differences between b=0 images in each encoding direction can occur due to subject movement.

21 All steps applied to images in the AP direction are also applied to images in the PA direction.

22 The fieldmaps will be used to correct both fMRI and dMRI data.

23 After making sure that the number of slices in the Z direction is a multiple of topup’s sub-sampling level (See https://fsl.fmrib.ox.ac.uk/fsl/fslwiki/topup/TopupUsersGuide#A--subsamp), defined in topup configuration file, by removing the appropriate number of slices from the top of the image.

24 Model 2: Deconvolution with sticks and a range of diffusivities

25 The reasoning behind these dimensionalities is: - 25 results in large scale network decomposition which matches much of the canonical RSN literature (Smith et al., 2009). - 100 corresponds to a more finely detailed parcellation. We found empirically that it was not useful to go even higher because, with volumetrically aligned data, raising the number of components did not significantly raise the number of non-artefact group level components (as opposed to using surface based analysis where the number of well-aligned small components can be much higher (Smith et al., 2013)

26 http://biobank.ctsu.ox.ac.uk/crystal/refer.cgi?id=9028.

27 http://fsl.fmrib.ox.ac.uk/fsl/fslwiki/FSLNets.

28 Both mixed-effects and fixed-effects generate huge z-statistics with this many subjects, so we are choosing here to simply work with the “average” activation given by using fixed-effects.

29 By testing on a reduced version of the final training set.

30 As all the cases flagged as “problems” would need a posterior visual inspection.

31 Age, age^2^, sex, age *×* sex, age^2^ *×* sex, average head motion during tfMRI, average head motion during rfMRI and head size

32 The proportion of the classes to classify is maintained in the folds, which is important in unbalanced situations

## References

Abe, S., Irimia, A., Van Horn, J. D., 2015. Quality Control Considerations for the E_ective Integration of Neuroimaging Data. In: Data Integration in the Life Sciences. Springer, pp. 195–201.

Afyouni, S., Nichols, T. E., 2017. Insight and inference for DVARS. bioRxiv, 125021.

Andersson, J. L., Graham, M. S., Zsoldos, E., Sotiropoulos, S. N., 2016. Incorporating outlier detection and replacement into a non-parametric framework for movement and distortion correction of diffusion mr images. NeuroImage 141, 556–572.

Andersson, J. L., Jenkinson, M., Smith, S., 2007a. Non-linear registration aka Spatial normalisation. Internal Technical Report TR07JA2, Oxford Centre for Functional Magnetic Resonance Imaging of the Brain, Department of Clinical Neurology, Oxford University, Oxford, UK, available at www.fmrib.ox.ac.uk/analysis/techrep for downloading.

Andersson, J. L., Skare, S., 2010. Image distortion and its correction in diffusion MRI. In: Jones, D. (Ed.), Diffusion MRI: theory, methods, and applications. Oxford University Press, Oxford, pp. 285–302.

Andersson, J. L., Skare, S., Ashburner, J., 2003. How to correct susceptibility distortions in spin-echo echo-planar images: application to diffusion tensor imaging. Neuroimage 20 (2), 870–888.

Andersson, J. L., Smith, S., Jenkinson, M., 2007b. Non-Linear Optimisation. Internal Technical Report TR07JA1, Oxford Centre for Functional Magnetic Resonance Imaging of the Brain, Department of Clinical Neurology, Oxford University, Oxford, UK, available at www.fmrib.ox.ac.uk/analysis/techrep for downloading.

Andersson, J. L., Sotiropoulos, S. N., 2015. Non-parametric representation and prediction of single-and multi-shell diffusion-weighted MRI data using Gaussian processes. NeuroImage 122, 166–176.

Andersson, J. L., Sotiropoulos, S. N., 2016. An integrated approach to correction for off-resonance effects and subject movement in diffusion MR imaging. NeuroImage 125, 1063–1078.

Baker, A. P., Brookes, M. J., Rezek, I. A., Smith, S. M., Behrens, T., Smith, P. J. P., Woolrich, M., 2014. Fast transient networks in spontaneous human brain activity. Elife 3, e01867.

Barch, D. M., Burgess, G. C., Harms, M. P., Petersen, S. E., Schlaggar, B. L., Corbetta, M., Glasser, M. F., Curtiss, S., Dixit, S., Feldt, C., et al., 2013. Function in the human connectome: task-fMRI and individual differences in behavior. Neuroimage 80, 169–189.

Basser, P. J., Mattiello, J., LeBihan, D., 1994. Estimation of the effective self-diffusion tensor from the NMR spin echo. Journal of Magnetic Resonance, Series B 103 (3), 247–254.

Beckmann, C., Smith, S., 2004. Probabilistic Independent Component Analysis for Functional Magnetic Resonance Imaging. IEEE Trans. on Medical Imaging 23 (2), 137–152.

Behrens, T., Berg, H. J., Jbabdi, S., Rushworth, M., Woolrich, M., 2007. Probabilistic diffusion tractography with multiple fibre orientations: What can we gain? Neuroimage 34 (1), 144–155.

Behrens, T., Woolrich, M., Jenkinson, M., Johansen-Berg, H., Nunes, R., Clare, S., Matthews, P., Brady, J., Smith, S., 2003. Characterization and propagation of uncertainty in diffusion-weighted MR imaging. Magnetic resonance in medicine 50 (5), 1077–1088.

Bennett, C. M., Miller, M. B., 2010. How reliable are the results from functional magnetic resonance imaging? Annals of the New York Academy of Sciences 1191 (1), 133–155.

Biobank, 2014. Biobank protocol specifications. http://biobank.ctsu.ox.ac.uk/crystal/docs/bmri_V4_23092014.pdf, accessed: 2017-02-01.

Biobank, 2016. Biobank First Public Imaging Release. http://biobank.ctsu.ox.ac.uk/crystal/label.cgi, accessed: 2016-01-01.

Biobank, 2017. New data from brain imaging and on heart attacks and strokes available. https://www.ukbiobank.ac.uk/2017/02/ new-data-from-brain-imaging-and-on-heart-attacks-and-strokes-available/, accessed: 2016-02-13.

Bouckaert, R. R., 2008. Bayesian Network Classifiers in Weka. http://wwww.cs.waikato.ac.nz/∼remco/weka.bn.pdf, accessed: 2016-09-7.

Breteler, M. M., Stöocker, T., Pracht, E., Brenner, D., Stirnberg, R., 2014. MRI in the Thineland study: A novel protocol for population neuroimaging. Alzheimer’s & Dementia: The Journal of the Alzheimer’s Association 10 (4), P92.

CBS, 2016. Qualitative Quality Control Manual. http://cbs.fas.harvard.edu/usr/mcmains/CBS_MRI_Qualitative_Quality_Control_Manual.pdf, accessed: 2016-04-22.

Chen, J., Liu, J., Calhoun, V. D., Arias-Vasquez, A., Zwiers, M. P., Gupta, C. N., Franke, B., Turner, J. A., 2014. Exploration of scanning effects in multi-site structural MRI studies. Journal of neuroscience methods 230, 37–50.

Cook, P., Bai, Y., Nedjati-Gilani, S., Seunarine, K., Hall, M., Parker, G., Alexander, D., 2006. Camino: open-source diffusion-MRI reconstruction and processing. In: 14th scientific meeting of the international society for magnetic resonance in medicine. Vol. 2759. Seattle WA, USA, p. 2759.

Craddock, C., Giavasis, S., Clark, D., Shezhad, Z., Pellman, J., Gorgolewski, C. F., Moodie, C. A., Esteban, O., 2016. PCP: Quality Assesment Protocol. http://preprocessed-connectomes-project.org/quality-assessment-protocol/, accessed: 2016-08-20.

Daducci, A., Canales-Rodríguez, E. J., Zhang, H., Dyrby, T. B., Alexander, D. C., Thiran, J.-P., 2015. Accelerated Microstructure Imaging via Convex Optimization (AMICO) from diffusion MRI data. NeuroImage 105, 32–44.

Dale, A. M., Fischl, B., Sereno, M. I., 1999. Cortical surface-based analysis I: Segmentation and surface reconstruction. Neuroimage 9, 179–94.

de Groot, M., Vernooij, M. W., Klein, S., Ikram, M. A., Vos, F. M., Smith, S., Niessen, W. J., Andersson, J. L., 2013. Improving alignment in tract-based spatial statistics: evaluation and optimization of image registration. Neuroimage 76, 400–411.

Deshmukh, M., Bhosale, U., 2010. Image fusion and image quality assessment of fused images. International Journal of Image Processing (IJIP) 4 (5), 484–508.

Domingos, P., 1999. Metacost: A general method for making classifiers cost-sensitive. In: Proceedings of the fifth ACM SIGKDD international conference on Knowledge discovery and data mining. ACM, pp. 155–164.

Duff, E. P., Vennart, W., Wise, R. G., Howard, M. A., Harris, R. E., Lee, M., Wartolowska, K., Wanigasekera, V., Wilson, F. J., Whitlock, M., et al., 2015. Learning to identify CNS drug action and efficacy using multistudy fmri data. Science translational medicine 7 (274), 274ra16–274ra16.

Duyn, J., 2013. Mr susceptibility imaging. Journal of magnetic resonance 229, 198–207.

Esteban, O., Birman, D., Schaer, M., Koyejo, O. O., Poldrack, R. A., Gorgolewski, K. J., 2017. Mriqc: Predicting quality in manual mri assessment protocols using no-reference image quality measures. bioRxiv, 111294.

Esteban, O., Gorgolewski, K. F., A., M. C., Tripplett, W., 2016. MRIQC: Image quality metrics for quality assessment of MRI. http://mriqc.readthedocs.io/en/latest/, accessed: 2016-08-20.

Filippini, N., MacIntosh, B., Hough, M., Goodwin, G., Frisoni, G., Smith, S., Matthews, P., Beckmann, C., Mackay, C., 2009. Distinct Patterns of Brain Activity in Young Carriers of the APOE-e4 Allele. Proc Natl Acad Sci USA (PNAS) 106, 7209–7214.

Focke, N. K., Helms, G., Kaspar, S., Diederich, C., Tóth, V., Dechent, P., Mohr, A., Paulus, W., 2011. Multi-site voxel-based morphometry-not quite there yet. Neuroimage 56 (3), 1164–1170.

Fréenay, B., Verleysen, M., 2014. Classification in the presence of label noise: a survey. IEEE transactions on neural networks and learning systems 25 (5), 845–869.

Friedman, L., Glover, G. H., 2006. Report on a multicenter fMRI quality assurance protocol. Journal of Magnetic Resonance Imaging 23 (6), 827–839.

Glasser, M. F., Sotiropoulos, S. N., Wilson, J. A., Coalson, T. S., Fischl, B., Andersson, J. L., Xu, J., Jbabdi, S., Webster, M., Polimeni, J. R., Van Essen, D. C., Jenkinson, M., 2013. The minimal preprocessing pipelines for the Human Connectome Project. NeuroImage 80, 105–24.

Glatard, T., Lewis, L. B., Ferreira da Silva, R., Adalat, R., Beck, N., Lepage, C., Rioux, P., Rousseau, M.-E., Sherif, T., Deelman, E., et al., 2015. Reproducibility of neuroimaging analyses across operating systems. Frontiers in neuroinformatics 9, 12.

Glover, G. H., Mueller, B. A., Turner, J. A., Van Erp, T. G., Liu, T. T., Greve, D. N., Voyvodic, J. T., Rasmussen, J., Brown, G. G., Keator, D. B., et al., 2012. Function biomedical informatics research network recommendations for prospective multicenter functional MRI studies. Journal of Magnetic Resonance Imaging 36 (1), 39–54.

GNC, 2014. The German National Cohort: aims, study design and organization. European Journal of Epidemiology 29 (5), 371.

Gorgolewski, K. J., Auer, T., Calhoun, V. D., Craddock, R. C., Das, S., Duff, E. P., Flandin, G., Ghosh, S. S., Glatard, T., Halchenko, Y. O., et al., 2016. The brain imaging data structure, a format for organizing and describing outputs of neuroimaging experiments. Scientific Data 3, 160044.

Greve, D., Fischl, B., 2009. “Accurate and robust brain image alignment using boundary-based registration”. NeuroImage 48, 63–72.

Greve, D. N., Mueller, B. A., Liu, T., Turner, J. A., Voyvodic, J., Yetter, E., Diaz, M., McCarthy, G., Wallace, S., Roach, B. J., et al., 2011. A novel method for quantifying scanner instability in fMRI. Magnetic resonance in medicine 65 (4), 1053–1061.

Griffanti, L., Douaud, G., Bijsterbosch, J., Evangelisti, S., Alfaro-Almagro, F., Glasser, M. F., Duff, E. P., Fitzgibbon, S., Westphal, R., Carone, D., et al., 2017 Hand classification of fmri ica noise components. NeuroImage 154, 188–205.

Griffanti, L., Salimi-Khorshidi, G., Beckmann, C., Auerbach, E., Douaud, G., Sexton, C., Zsoldos, E., Ebmeier, K., Filippini, N., Mackay, C., Moeller, S., Xu, J., Yacoub, E., Baselli, G., Ugurbil, K., Miller, K., Smith, S., 2014. ICA-based artefact removal and accelerated fMRI acquisition for improved Resting State Network imaging. NeuroImage 95, 232–247.

Griffanti, L., Zamboni, G., Khan, A., Li, L., Bonifacio, G., Sundaresan, V., Schulz, U. G., Kuker, W., Battaglini, M., Rothwell, P. M., et al., 2016. BIANCA (Brain Intensity AbNormality Classification Algorithm): A new tool for automated segmentation of white matter hyperintensities. NeuroImage 141, 191–205.

Gronenschild, E. H., Habets, P., Jacobs, H. I., Mengelers, R., Rozendaal, N., Van Os, J., Marcelis, M., 2012. The effects of freesurfer version, workstation type, and macintosh operating system version on anatomical volume and cortical thickness measurements. PloS one 7 (6), e38234.

Gunstad, J., Cohen, R. A., Tate, D. F., Paul, R. H., Poppas, A., Hoth, K., Macgregor, K. L., Jefferson, A. L., 2005. Blood pressure variability and white matter hyperintensities in older adults with cardiovascular disease. Blood pressure 14 (6), 353–358.

Haacke, E., Xu, Y., Cheng, Y., Reichenbach, J., 2004. Susceptibility-weighted imaging (SWI). Magnetic Resonance in Medicine 52, 612–618.

Hall, M., Frank, E., Holmes, G., Pfahringer, B., Reutemann, P., Witten, I. H., 2009. The weka data mining software: an update. ACM SIGKDD explorations newsletter 11 (1), 10–18.

Hariri, A. R., Tessitore, A., Mattay, V. S., Fera, F., Weinberger, D. R., 2002. The amygdala response to emotional stimuli: a comparison of faces and scenes. Neuroimage 17 (1), 317–323.

Hasan, K. M., 2007. A framework for quality control and parameter optimization in diffusion tensor imaging: theoretical analysis and validation. Magnetic resonance imaging 25 (8), 1196–1202.

Hernández, M., Guerrero, G. D., Cecilia, J. M., García, J. M., Inuggi, A., Jbabdi, S., Behrens, T. E., Sotiropoulos, S. N., 2013. Accelerating fibre orientation estimation from diffusion weighted magnetic resonance imaging using GPUs. PloS one 8 (4), e61892.

Hyvärinen, A., 1999. Fast and robust fixed-point algorithms for independent component analysis. IEEE Transactions on Neural Networks 10 (3), 626–634.

Jahanshad, N., Kochunov, P. V., Sprooten, E., Mandl, R. C., Nichols, T. E., Almasy, L., Blangero, J., Brouwer, R. M., Curran, J. E., de Zubicaray, G. I., et al., 2013. Multi-site genetic analysis of diffusion images and voxelwise heritability analysis: A pilot project of the ENIGMA-DTI working group. Neuroimage 81, 455–469.

Jbabdi, S., Sotiropoulos, S. N., Savio, A. M., Gra∼na, M., Behrens, T. E., 2012. Model-based analysis of multishell diffusion MR data for tractography: How to get over _tting problems. Magnetic Resonance in Medicine 68 (6), 1846–1855.

Jenkinson, M., Bannister, P., Brady, J., Smith, S., 2002. Improved Optimisation for the Robust and Accurate Linear Registration and Motion Correction of Brain Images. NeuroImage 17 (2), 825–841.

Jenkinson, M., Beckmann, C. F., Behrens, T. E., Woolrich, M. W., Smith, S., 2012. FSL. Neuroimage 62 (2), 782–790.

Jenkinson, M., Smith, S., 2001. A Global Optimisation Method for Robust Affine Registration of Brain Images. Medical Image Analysis 5 (2), 143–156.

Jezzard, P., Balaban, R. S., 1995. Correction for geometric distortion in echo planar images from B0 field variations. Magnetic resonance in medicine 34 (1), 65–73.

John, G. H., Langley, P., 1995. Estimating continuous distributions in bayesian classifiers. In: Proceedings of the Eleventh conference on Uncertainty in artificial intelligence. Morgan Kaufmann Publishers Inc., pp. 338–345.

Kittler, J., Hatef, M., Duin, R. P., Matas, J., 1998. On combining classifiers. IEEE transactions on pattern analysis and machine intelligence 20 (3), 226–239.

Kuncheva, L. I., 2004. Combining pattern classifiers: methods and algorithms. John Wiley & Sons.

Li, X., Morgan, P. S., Ashburner, J., Smith, J., Rorden, C., 2016. The first step for neuroimaging data analysis: DICOM to NIfTI conversion. Journal of Neuroscience Methods 264, 47–56.

Liao, D., Cooper, L., Cai, J., Toole, J. F., Bryan, N. R., Hutchinson, R. G., Tyroler, H. A., 1996. Presence and severity of cerebral white matter lesions and hypertension, its treatment, and its control. Stroke 27 (12), 2262–2270.

Liu, Z., Wang, Y., Gerig, G., Gouttard, S., Tao, R., Fletcher, T., Styner, M., 2010. Quality control of diffusion weighted images. In: SPIE medical imaging. International Society for Optics and Photonics, pp. 76280J–76280J.

Marcus, D. S., Harms, M. P., Snyder, A. Z., Jenkinson, M., Wilson, J. A., Glasser, M. F., Barch, D. M., Archie, K. A., Burgess, G. C., Ramaratnam, M., et al., 2013. Human Connectome Project informatics: quality control, database services, and data visualization. Neuroimage 80, 202–219.

Miller, K. L., Alfaro-Almagro, F., Bangerter, N. K., Thomas, D. L., Yacoub, E., Xu, J., Bartsch, A. J., Jbabdi, S., Sotiropoulos, S. N., Andersson, J. L., Griffanti, L., Douaud, G., Okell, T. W., Weale, P., Dragonu, I., Garratt, S., Hudson, S., Collins, R., Jenkinson, M., Matthews, P. M., Smith, S., 2016. Multimodal population brain imaging in the UK Biobank prospective epidemiological study. Neuroimage.

Moeller, S., Yacoub, E., Olman, C. A., Auerbach, E., Strupp, J., Harel, N., Uğurbil, K., 2010. Multiband multislice ge-epi at 7 tesla, with 16-fold acceleration using partial parallel imaging with application to high spatial and temporal whole-brain fmri. Magnetic Resonance in Medicine 63 (5), 1144–1153.

Moorthy, A. K., Bovik, A. C., 2010. A two-step framework for constructing blind image quality indices. Signal Processing Letters, IEEE 17 (5), 513–516.

Mori, S., Wakana, S., Van Zijl, P. C., Nagae-Poetscher, L., 2005. MRI atlas of human white matter. Vol. 16. Am Soc Neuroradiology.

Mortamet, B., Bernstein, M. A., Jack, C. R., Gunter, J. L., Ward, C., Britson, P. J., Meuli, R., Thiran, J.-P., Krueger, G., 2009. Automatic Quality Assessment in structural brain Magnetic Resonance Imaging. Magnetic Resonance in Medicine 62 (2), 365–372.

Mugler, J. P., 2014. Optimized three-dimensional fast-spin-echo mri. Journal of Magnetic Resonance Imaging 39 (4), 745–767.

Mugler, J. P., Brookeman, J. R., 1990. Three-dimensional magnetization-prepared rapid gradient-echo imaging (3d mp rage). Magnetic Resonance in Medicine 15 (1), 152–157.

Nichols, T., 2013. Notes on creating a standardized version of DVARS.

Patenaude, B., Smith, S., Kennedy, D., Jenkinson, M., 2011. A Bayesian Model of Shape and Appearance for Subcortical Brain Segmentation. NeuroImage 56 (3), 907–922.

Power, J. D., Barnes, K. A., Snyder, A. Z., Schlaggar, B. L., Petersen, S. E., 2012. Spurious but systematic correlations in functional connectivity MRI networks arise from subject motion. Neuroimage 59 (3), 2142–2154.

Robinson, E. C., Jbabdi, S., Glasser, M. F., Andersson, J. L., Burgess, G. C., Harms, M. P., Smith, S., Van Essen, D. C., Jenkinson, M., 2014. Msm: A new flexible framework for multimodal surface matching. Neuroimage 100, 414–426.

Salimi-Khorshidi, G., Douaud, G., Beckmann, C., Glasser, M., Griffanti, L., Smith, S., 2014. Automatic Denoising of Functional MRI Data: Combining Independent Component Analysis and Hierarchical Fusion of classifiers. NeuroImage 90, 449–468.

Schram, M. T., Sep, S. J., van der Kallen, C. J., Dagnelie, P. C., Koster, A., Schaper, N., Henry, R. M., Stehouwer, C. D., 2014. The Maastricht Study: an extensive phenotyping study on determinants of type 2 diabetes, its complications and its comorbidities. European journal of epidemiology 29 (6), 439–451.

Smith, S., November 2002. Fast Robust Automated Brain Extraction. Human Brain Mapping 17 (3), 143–155.

Smith, S., 2012. The Future of FMRI Connectivity. NeuroImage 62, 1257–1266.

Smith, S., Beckmann, C. F., Andersson, J. L., Auerbach, E. J., Bijsterbosch, J., Douaud, G., Duff, E., Feinberg, D. A., Griffanti, L., Harms, M. P., et al., 2013. Resting-state fMRI in the human connectome project. Neuroimage 80, 144–168.

Smith, S., Fox, P. T., Miller, K. L., Glahn, D. C., Fox, P. M., Mackay, C. E., Filippini, N., Watkins, K. E., Toro, R., Laird, A. R., et al., 2009. Correspondence of the brain’s functional architecture during activation and rest. Proceedings of the National Academy of Sciences 106 (31), 13040–13045.

Smith, S., Hyvärinen, A., Varoquaux, G., Miller, K., Beckmann, C., 2014. Group-PCA for very large fMRI datasets. NeuroImage 101, 738–749.

Smith, S., Jenkinson, M., Johansen-Berg, H., Rueckert, D., Nichols, T. E., Mackay, C. E., Watkins, K. E., Ciccarelli, O., Cader, M. Z., Matthews, P. M., et al., 2006. Tract-based spatial statistics: voxelwise analysis of multi-subject diffusion data. Neuroimage 31 (4), 1487–1505.

Smith, S., Zhang, Y., Jenkinson, M., Chen, J., Matthews, P., Federico, A., De Stefano, N., 2002. Accurate, Robust and Automated Longitudinal and Cross-Sectional Brain Change Analysis. NeuroImage 17 (1), 479–489.

Song, X.-W., Dong, Z.-Y., Long, X.-Y., Li, S.-F., Zuo, X.-N., Zhu, C.-Z., He, Y., Yan, C.-G., Zang, Y.-F., 2011. REST: a toolkit for resting-state functional magnetic resonance imaging data processing. PloS one 6 (9), e25031.

Sotiropoulos, S., Moeller, S., Jbabdi, S., Xu, J., Andersson, J., Auerbach, E., Yacoub, E., Feinberg, D., Setsompop, K., Wald, L., et al., 2013a. Effects of image reconstruction on fiber orientation mapping from multichannel diffusion MRI: reducing the noise floor using SENSE. Magnetic resonance in medicine 70 (6), 1682–1689.

Sotiropoulos, S. N., Jbabdi, S., Xu, J., Andersson, J. L., Moeller, S., Auerbach, E. J., Glasser, M. F., Hernandez, M., Sapiro, G., Jenkinson, M., et al., 2013b. Advances in diffusion MRI acquisition and processing in the Human Connectome Project. Neuroimage 80, 125–143.

Sudlow, C., Gallacher, J., Allen, N., Beral, V., Burton, P., Danesh, J., Downey, P., Elliott, P., Green, J., Landray, M., et al., 2015. UK Biobank: an Open Access resource for identifying the causes of a wide range of complex diseases of middle and old age. PLoS medicine 12 (3), 1–10.

Vidaurre, D., Quinn, A. J., Baker, A. P., Dupret, D., Tejero-Cantero, A., Woolrich, M. W., 2016. Spectrally resolved fast transient brain states in electrophysiological data. Neuroimage 126, 81–95.

Wakana, S., Caprihan, A., Panzenboeck, M. M., Fallon, J. H., Perry, M., Gollub, R. L., Hua, K., Zhang, J., Jiang, H., Dubey, P., et al., 2007. Reproducibility of quantitative tractography methods applied to cerebral white matter. Neuroimage 36 (3), 630–644.

Wei, X., Warfield, S. K., Zou, K. H., Wu, Y., Li, X., Guimond, A., Mugler, J. P., Benson, R. R., Wolfson, L., Weiner, H. L., et al., 2002. Quantitative analysis of MRI signal abnormalities of brain white matter with high reproducibility and accuracy. Journal of Magnetic Resonance Imaging 15 (2), 203–209.

Woodard, J. P., Carley-Spencer, M. P., 2006. No-reference image quality metrics for structural MRI. Neuroinformatics 4 (3), 243–262.

Woolrich, M., Ripley, B., Brady, J., Smith, S., 2001. Temporal Autocorrelation in Univariate Linear Modelling of FMRI Data. NeuroImage 14 (6), 1370–1386.

Xu, J., Moeller, S., Auerbach, E. J., Strupp, J., Smith, S. M., Feinberg, D. A., Yacoub, E., Uğurbil, K., 2013. Evaluation of slice accelerations using multiband echo planar imaging at 3t. Neuroimage 83, 991–1001.

Zhang, H., Schneider, T., Wheeler-Kingshott, C. A., Alexander, D. C., 2012. NODDI: practical in vivo neurite orientation dispersion and density imaging of the human brain. Neuroimage 61 (4), 1000–1016.

Zhang, Y., Brady, M., Smith, S., 2001. Segmentation of Brain MR Images through a Hidden Markov Random Field Model and the Expectation Maximization Algorithm. IEEE Trans. on Medical Imaging 20 (1), 45–57.

